# A Spatially Resolved Atlas of Alternative Polyadenylation Across 18 Human Tissues and 76 Disease States

**DOI:** 10.1101/2025.08.27.672753

**Authors:** Zehang Jiang, Zhuochao Min, Zhanying Wu, Yubin Chen, Zhiyong Wu, Huashu Wen, Cheng Wu, Jia Guo, Ke Si, Douyue Li, Guoying Wang, Shuai Mao, Weizhong Li, Binghui Zeng, Wenliang Zhang

## Abstract

Alternative polyadenylation (APA) is a key regulator of gene expression and cellular dynamics, yet systematic investigations of spatially resolved APA across diverse human tissues remain limited. Here, we developed SpatialAPA (https://github.com/Omicslab-Zhang/spatialAPA), a framework that benchmarks multiple APA identification methods and integrates spatial APA data with gene expression and cellular dynamics at spatial resolution. Applying SpatialAPA to 363 spatial transcriptomic data from 56 projects across 18 human tissues and 76 diseases, we constructed a spatially resolved APA atlas comprising 346,932 APA events across 52,175 genes. This atlas reveals organ–specific APA patterns and provides new insights into how APA regulates tissue homeostasis and disease progression beyond transcriptional control. To ensure cross–sample comparability, we applied batch correction, while spatial cell deconvolution uncovered cell–type–specific dynamics and interactions. In triple–negative breast cancer, integrated spatial and single–cell analyses identified *TSPAN8*–positive epithelial subpopulations whose distinct APA regulation and transcriptional programs drive differentiation and malignant progression. To facilitate community access, we developed an online platform (http://www.biomedical-web.com/spatialAPAdb/home) for exploring APA, gene expression, and cellular dynamics in health and disease. Together, this study establishes the first comprehensive spatial APA atlas, providing a valuable resource and analytical framework for investigating molecular mechanisms and therapeutic targets.

## Introduction

Alternative polyadenylation (APA) is a critical post–transcriptional regulatory mechanism that generates mRNA isoforms with distinct 3’ untranslated regions (UTRs) by adding poly(A) tails at various sites [1, 2]. This process profoundly influences mRNA stability, translation efficiency, and cellular localization, ultimately affecting gene expression and cellular function [3, 4]. APA occurs widely across tissues, cell types, and developmental stages, playing vital roles in cellular processes and being closely linked to a variety of disease states [3, 5, 6]. While the regulatory significance of APA is well–established, its spatial dynamics within human tissues, particularly in physiological and pathological contexts, remain largely unexplored. Understanding these spatial dynamics is essential for uncovering tissue–specific regulatory mechanisms and disease–related alterations, which may lead to the discovery of novel biomarkers and therapeutic targets.

Existing resources have been developed to investigate APA events at the tissue level, providing valuable insights into the regulatory mechanisms of APA across species and disease contexts [7–14]. For instance, APAatlas [9] documents APA events in 53 normal human tissues, creating a comprehensive dataset of APA under physiological conditions. Despite these advancements, these existing APA resources primarily rely on bulk RNA sequencing data, focusing on tissue–level analyses while neglecting cell–specific and spatial distributions of APA, as well as the high–resolution dynamic changes of APA events within tissues. The emergence of single–cell RNA sequencing (scRNA–seq) has significantly enhanced our ability to analyze gene expression at single–cell resolution, enabling the identification of APA events across tissues, biological processes, and disease states [15]. Tools such as SCAPE, scUTRquant [16], SCAPE [17], scDAPA [18], Sierra [19], scAPAtrap [20], and scDaPars [21] have been crucial in identifying subpopulations and highlighting the role of APA in distinguishing cell types. Databases like scAPAatlas [22], scAPAdb [23], and scTEA–db [24] have provided novel APA events and terminal exons identified from human and mouse scRNA–seq data, offering valuable insights into their roles in cell identity and function. a major limitation of scRNA–seq is the dissociation of tissue samples, which eliminates the crucial spatial context needed to understand the regulatory roles of APA in intact tissues.

To overcome those limitations, spatial transcriptomics (ST) technologies [25], such as 10x Genomics Visium, have been developed to provide gene expression data with spatial resolution from tissue sections. While ST has successfully revealed spatial gene expression patterns in healthy and diseased tissues, most studies have focused primarily on gene expression at the tissue level [26–30], without fully utilizing the additional insights that spatial APA analysis can provide. Single–cell and spatial transcriptomic analyses share conceptual similarities, allowing strategies from single–cell APA identification to be applied in spatial contexts. For example, Kang et al. developed Infernape to uncover cell type–specific and spatially resolved APA in the brain [31], while stAPAminer, built on scAPAtrap, identifies spatial APA patterns in transcriptomic studies [32]. However, existing methods are largely limited to specific single–cell APA algorithms, restricting their generalizability. This highlights the need for a versatile spatial APA framework that can integrate multiple single–cell APA approaches, enabling tailored strategy selection and systematic benchmarking across diverse tissues to enhance reliability and applicability.

The growing availability of ST datasets presents unprecedented opportunities to study APA at spatial resolution. Effective spatial APA analysis, however, requires comprehensive preprocessing, including quality control, APA event detection, data integration, batch correction, and advanced analytical tools. Despite the wealth of data, dedicated methods to quantify and integrate spatial APA with gene expression and cellular dynamics remain scarce. Developing a comprehensive framework for spatial APA analysis is therefore essential to construct a spatially resolved APA atlas, reveal tissue–specific APA patterns, and provide insights into gene regulation, cell interactions, and disease mechanisms.

To address this challenge, we developed the SpatialAPA framework, which integrates multiple single–cell APA methods to analyze APA at cellular and spatial levels alongside gene expression and cellular dynamics. We further benchmarked and optimized these methods to enhance spatial APA identification, providing a robust foundation for systematic spatial APA studies. Using SpatialAPA, we systematically compiled available ST data to construct a spatially resolved APA atlas for human health and disease. This framework enables high–resolution mapping of APA usage across tissue regions, revealing spatial and cell type–specific regulation. The atlas uncovers how APA contributes to tissue–specific gene expression, cell interactions, and disease mechanisms, with particular relevance to cancer. By characterizing the spatial dynamics of APA, this resource advances our understanding of gene regulation in health and disease and provides a foundation for developing targeted therapies. Overall, this study bridges ST and APA research, offering a valuable platform for exploring disease mechanisms, identifying biomarkers, and guiding therapeutic interventions.

## Results

### A comprehensive analysis framework for analyzing and integrating spatial APA data with gene expression and cellular dynamics

Despite significant progress in ST for revealing spatial gene expression patterns in healthy and diseased tissues, most studies have been limited to gene expression analysis at the tissue level [26–30]. This is largely due to the absence of dedicated analysis frameworks for spatially resolved APA. As a result, the valuable APA information provided by ST sequencing has often been overlooked, hindering its integration with gene expression and cellular dynamics data. To address this gap, we developed SpatialAPA, a comprehensive analysis framework designed to analyze and integrate spatial APA data with gene expression and tissue cell dynamics (as shown in **Figure 1**).

**Figure 1.**
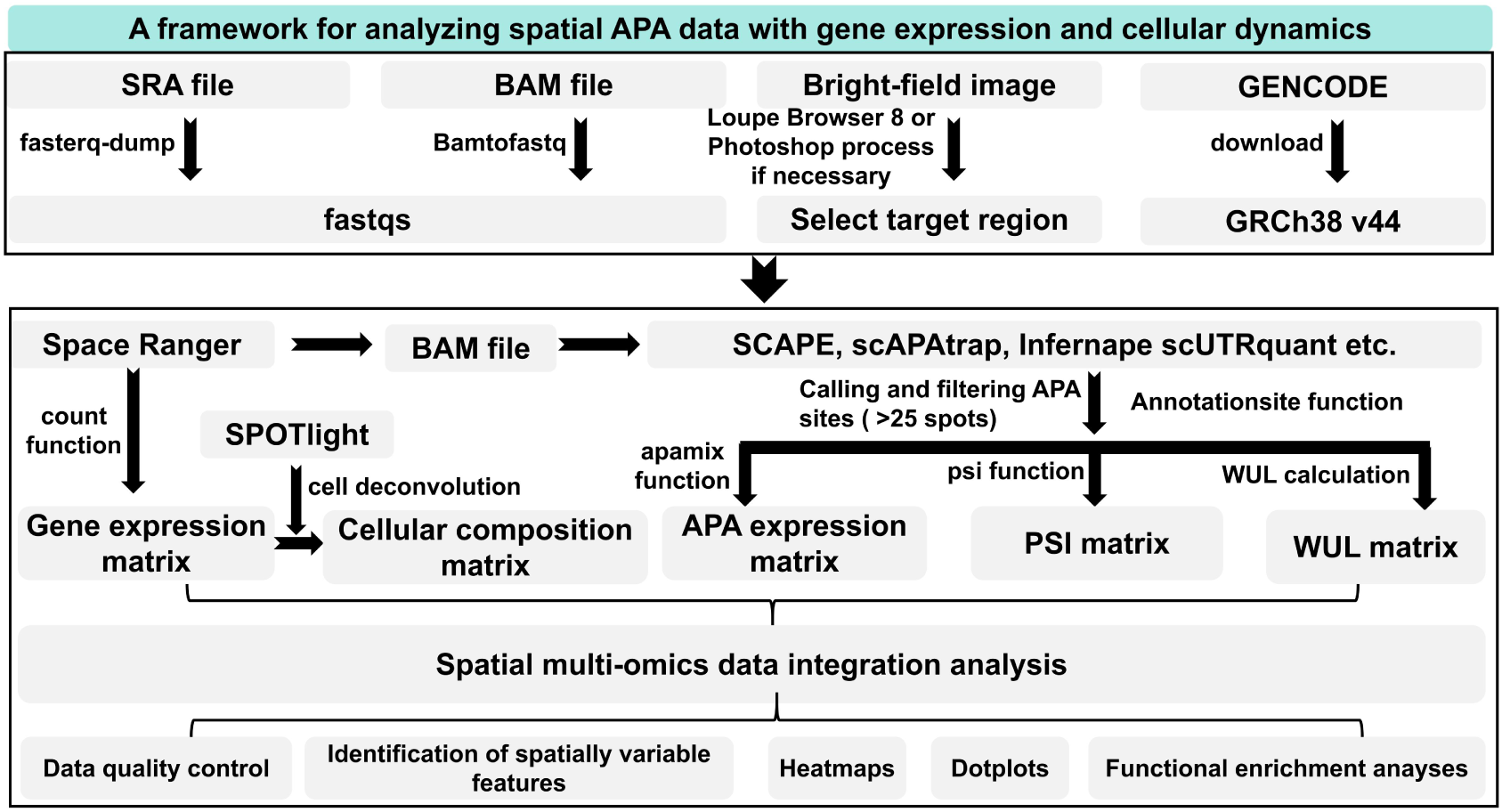
The SpatialAPA framework for analyzing spatial alternative polyadenylation (APA) data with gene expression and cellular dynamics. PSI (polyadenylation site usage index) quantifies the relative usage of alternative polyadenylation sites within a gene, reflecting APA site selection. WUL (weighted 3′ UTR length) measures the effective 3′ UTR length of a gene by weighting each isoform according to its expression level, thereby capturing overall APA-mediated changes in 3′ UTR length.

This framework is built on Snakemake and is compatible with Space Ranger v3.0.0 [33], Scanpy, SPOTlight [34], and several widely used single–cell APA detection algorithms (including SCAPE v1.0 [17], scAPAtrap v1.0 [20], scUTRquant v0.5.0 [16], and Infernape v1.0 [31], enabling end–to–end integrative analysis of spatial transcriptomics data (Figure 1). Specifically, the Space Ranger v3.0.0 [33] and SPOTlight 1.6.3 [31] are employed for spatial gene expression quantification and spatial cell deconvolution, respectively, thereby resolving cell–type distributions at spatial resolution. In addition, users can flexibly specify different single–cell APA algorithms for spatial APA identification and quantification. The SpatialAPA framework, fully described in the Methods section, is publicly available on GitHub (https://github.com/Omicslab-Zhang/spatialAPA) and National Genomics Data Center (NGDC) BioCode (https://ngdc.cncb.ac.cn/biocode/tool/7716).

Moreover, we evaluated thresholds of 0, 25, 50, and 100 spots to filter sparse APA events. The results showed that using a threshold of 25 spots not only substantially improved the accuracy of APA detection but also retained a sufficient number of APA events, indicating that this threshold achieves an optimal balance between detection precision and event coverage (**Figure 2A**). Given the potential positional shifts of APA sites across samples, we evaluated correction windows of 0, ±25, and ±50 nt. The results showed that a ±25 nt window markedly improved concordance with known APA sites, whereas extending the window to ±50 nt did not yield further benefits and could even introduce overcorrection (Figure 2B).

**Figure 2.**
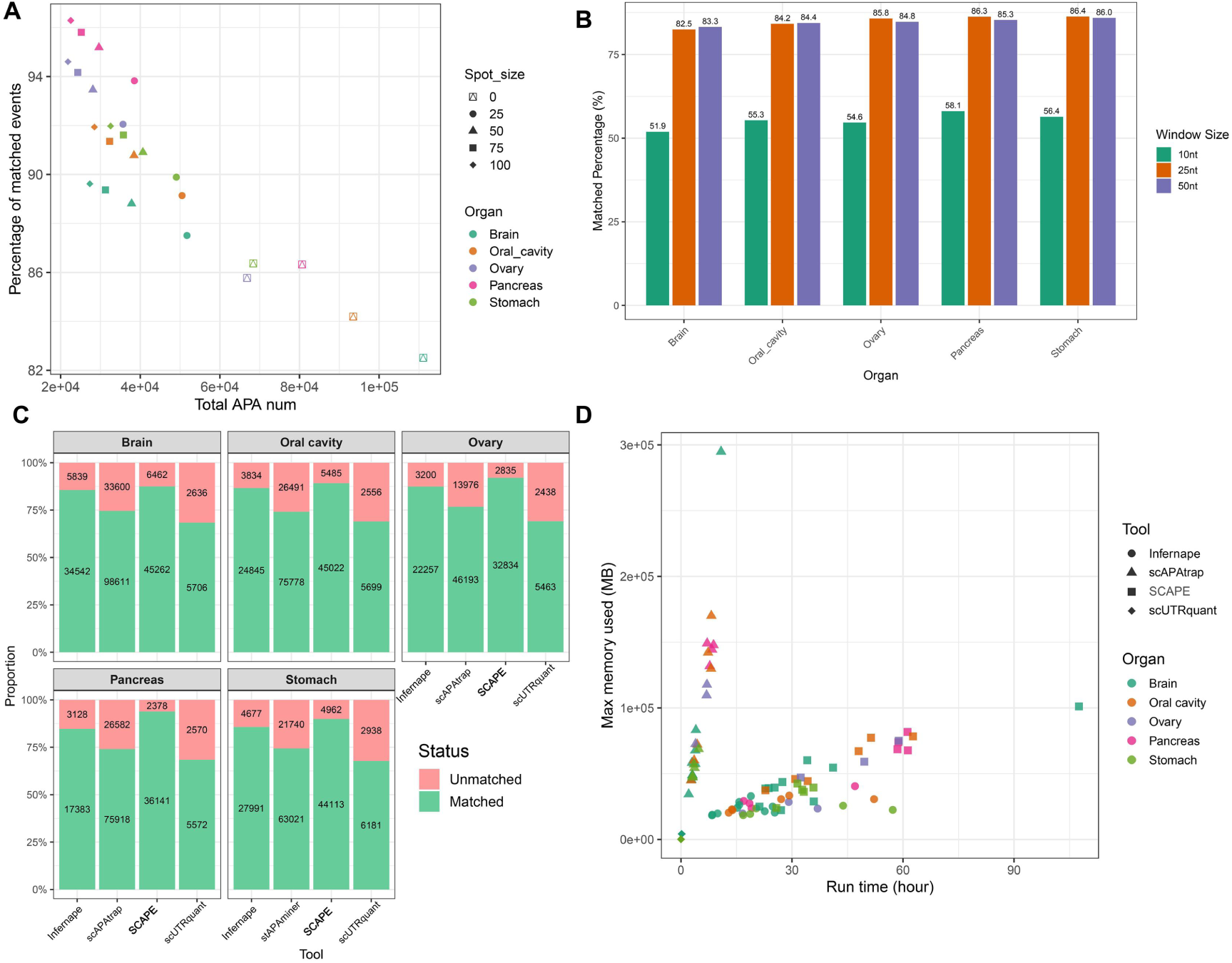
Benchmarking sparse APA filtering, shift correction, and different APA identification methods within the SpatialAPA framework. **(A)** Filtering of sparse APA sites based on the number of supporting spots. **(B)** Shift correction of APA site identification. **(C)** Comparison of different single-cell APA methods in terms of the number of identified APA sites and their consistency with known APA databases. **(D)** Comparison of peak memory usage and computation time across methods in 8 threads.

In terms of algorithmic performance and applicability, scAPAtrap v1.0 identified the largest number of APA sites but achieved only ∼75% concordance with known sites from human reference resources integrated from Gencode v44, PolyASite 3.0, and PolyA_DB (Figure 2C). In contrast, SCAPE v1.0, although detecting slightly fewer sites, showed the highest concordance (>90%) with these reference databases (Figure 2C). Regarding resource consumption (8 threads), scAPAtrap required a median of ∼100 GB memory compared with ∼50 GB for SCAPE v1.0, i.e., double the usage (Figure 2D). However, scAPAtrap v1.0 showed a modest advantage in runtime for individual samples (Figure 2D). scUTRquant v0.5.0 and Infernape v1.0 consumed fewer resources and completed more quickly, with moderate accuracy, but identified substantially fewer APA sites than SCAPE v1.0 and scAPAtrap (Figure 2C and 2D). Notably, these performance patterns were consistently observed across multiple tissues, including brain, oral cavity, ovary, pancreas, and stomach (Figure 2C and 2D). Overall, these results indicate that SCAPE v1.0 offers the best balance of accuracy, efficiency, and resource usage among the evaluated methods, making it the recommended choice for APA identification within the SpatialAPA framework.

### A spatially resolved atlas of alternative polyadenylation across 18 human tissue organs and 76 disease states

To construct a spatially resolved atlas of APA, we manually curated 56 publicly available human ST projects from the NCBI SRA database. These datasets cover 18 human tissue organs and 76 disease states, comprising 363 spatial sections and 804,276 transcriptomic spots (**Figure 3A**). The majority of data come from tissues associated with tumors, immune inflammation, and neurological diseases. The organs with the highest representation include skin (9 datasets), kidney (8 datasets), breast (7 datasets), liver (5 datasets), and ovary (5 datasets) (Figure S1A). The most extensively studied organs in terms of spatial sections are kidney (44 sections), brain (38 sections), skin (37 sections), breast (33 sections), and pancreas (31 sections) (Figure S1B). Common conditions represented in the dataset include breast carcinoma, ovarian carcinoma, lumbar spinal diseases, oral squamous cell carcinoma, atopic dermatitis, intraductal papillary mucinous neoplasm, and glioblastoma (Figure S1C). Furthermore, the atlas highlights the global contributions to ST data, with the United States, China, South Korea, the United Kingdom, and Russia being the primary contributors (Figure S1D).

**Figure 3.**
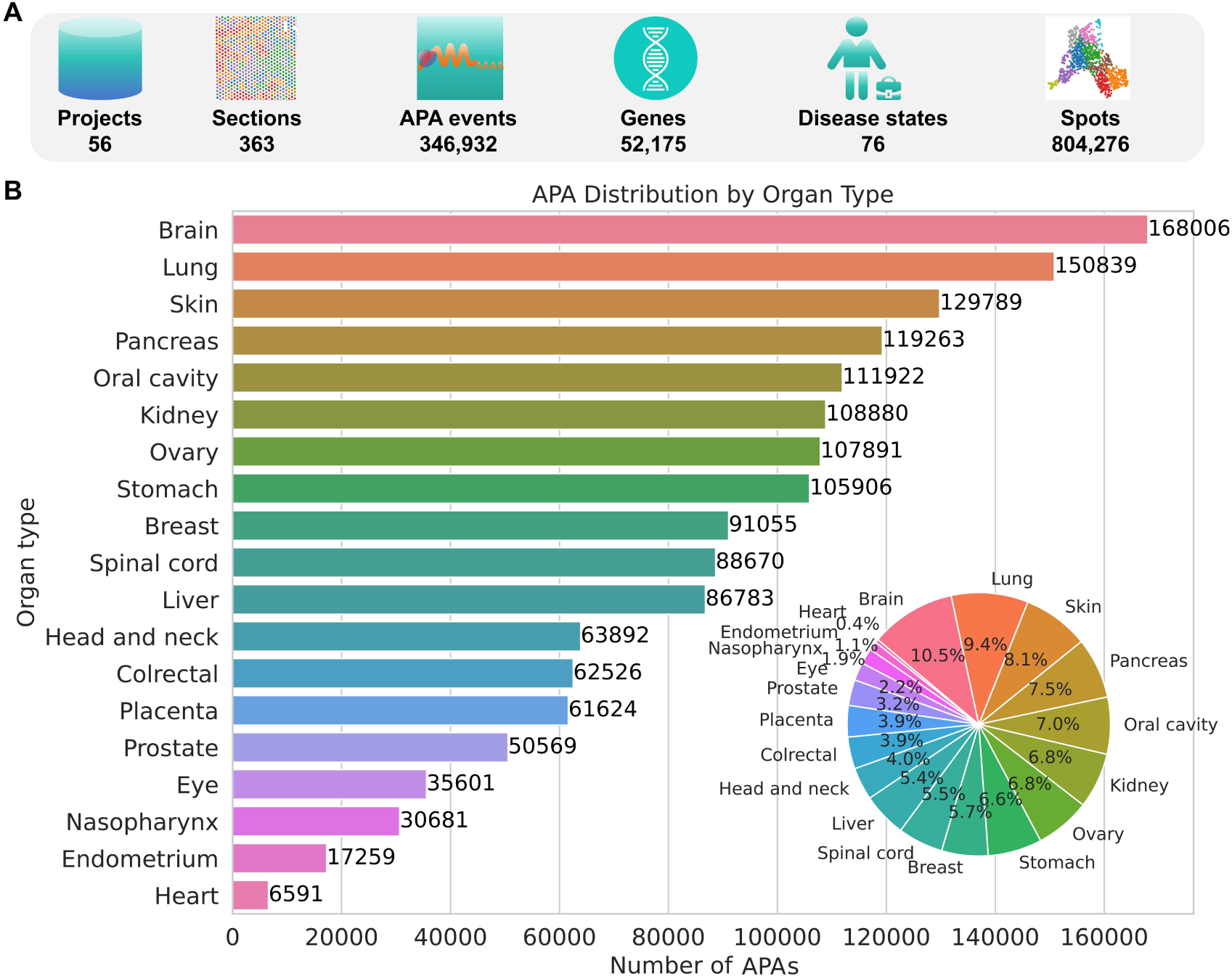
Overview of the spatially resolved APA atlas. **(A)** Data summary. **(B)** Distribution of APA events across human tissues.

Considering the significant advantages of SCAPE v1.0 in both the number and accuracy of spatial APA event detection, we implemented the SCAPE pipeline within the SpatialAPA framework to analyze all 363 spatial transcriptomic sections. This analysis identified 346,932 APA events across 52,175 genes, spanning 18 organs and 76 disease states (Figure 3A). The brain, lung, skin, pancreas, and oral cavity were the five organs with the highest number of APA events (Figure 3B). Further analysis revealed significant differences in sequencing data quantity, gene detection rates, APA event counts, and associated genes across different organ systems (**Figure S2A**). Despite the variability in sequencing data quantity, the patterns of detected genes, APA events, and WUL scores showed greater consistency across organs (**Figure S2A and S2B**).

At the genome–wide level, we observed significant organ–specific consistency in both APA events and gene expression patterns (**Figure 4A**), reinforcing the accuracy and reliability of this spatial APA data analysis framework. Compared to gene expression levels, APA events exhibited stronger organ–specificity, emphasizing the crucial regulatory role of APA in tissue homeostasis and disease progression (Figure 4B). These findings provide new insights into the complex, organ–specific dynamics of APA and its impact on health and disease, offering potential avenues for therapeutic exploration.

**Figure 4.**
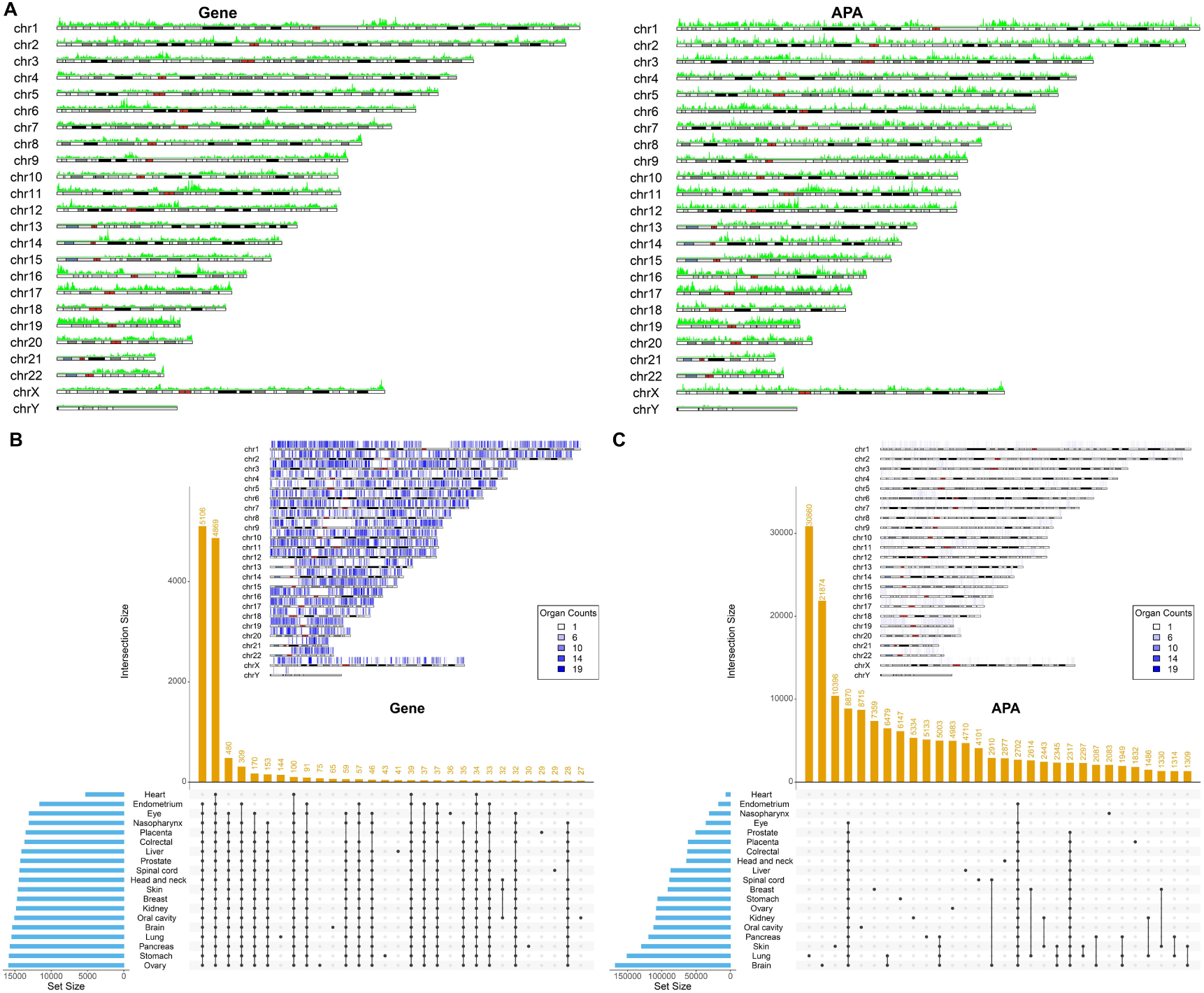
Genome-wide distribution and cross-tissue comparison of gene expression and APA events. **(A)** Distribution of gene expression levels and APA event abundance across chromosomes in 100 Mb windows. **(B)** Shared gene expression across different tissues and organs. **(C)** Shared APA events across different tissues and organs.]

Considering potential batch effects in ST data, independent dimensionality reduction and clustering of each spatial section could yield inconsistent results and hinder comparisons of APA events and gene expression dynamics. To address this, we grouped the 363 tissue sections into 18 organ categories and applied robust principal component analysis (RPCA) in Seurat v5.0.1 for integration and batch effect correction (see Materials and Methods). Before integration, UMAP analysis revealed pronounced batch effects among samples from the same organ (Figure S3A). After RPCA, these effects were substantially reduced, with clustering results across consecutive sections of the same tissue showing improved consistency and alignment with cellular deconvolution outcomes (Figure S3B). A similar improvement was observed in sections from different patients with the same pathology type (Figure S3C). Quantitatively, ARI (0.051) and Jaccard index (0.022) comparisons confirmed that RPCA integration effectively altered clustering structure and mitigated batch effects. Together, these results demonstrate that RPCA–based integration meaningfully reduces batch effects and ensures clustering consistency across spatial sections, thereby providing a robust foundation for comparative analyses of APA dynamics and gene expression across diverse physiological and pathological contexts.

### Spatial and single–cell analyses reveal unique APA dynamics in *TSPAN8*–positive TNBC epithelial cells

To validate the utility of the spatially resolved atlas, we conducted a study using a ST dataset of breast cancer (BC), which includes four major subtypes: HER2+ BC, triple–negative breast cancer (TNBC), Luminal – HER2+ BC, and Luminal BC. The analysis revealed significant heterogeneity in cell distribution across these BC subtypes (**Figure 5A** and Figure S4). In spots enriched for cancer epithelial cells, unlike HER2+ breast cancer, the proportion of cancer epithelial cells in TNBC, Luminal–HER2+, and Luminal breast cancer patients was significantly negatively correlated with the proportions of cancer associated fibroblasts (CAFs) and myeloid cells, and positively correlated with that of normal epithelial cells, suggesting that cancer epithelial cells may inhibit the infiltration of CAFs and myeloid cells while influencing the function of normal epithelial cells (Figure 5B). Moreover, in Luminal BC, the proportion of cancer epithelial cells was also significantly negatively correlated with the proportion of T cells (Figure 5B). These findings indicate that distinct molecular subtypes of breast cancer exhibit markedly different cellular dynamics.

**Figure 5.**
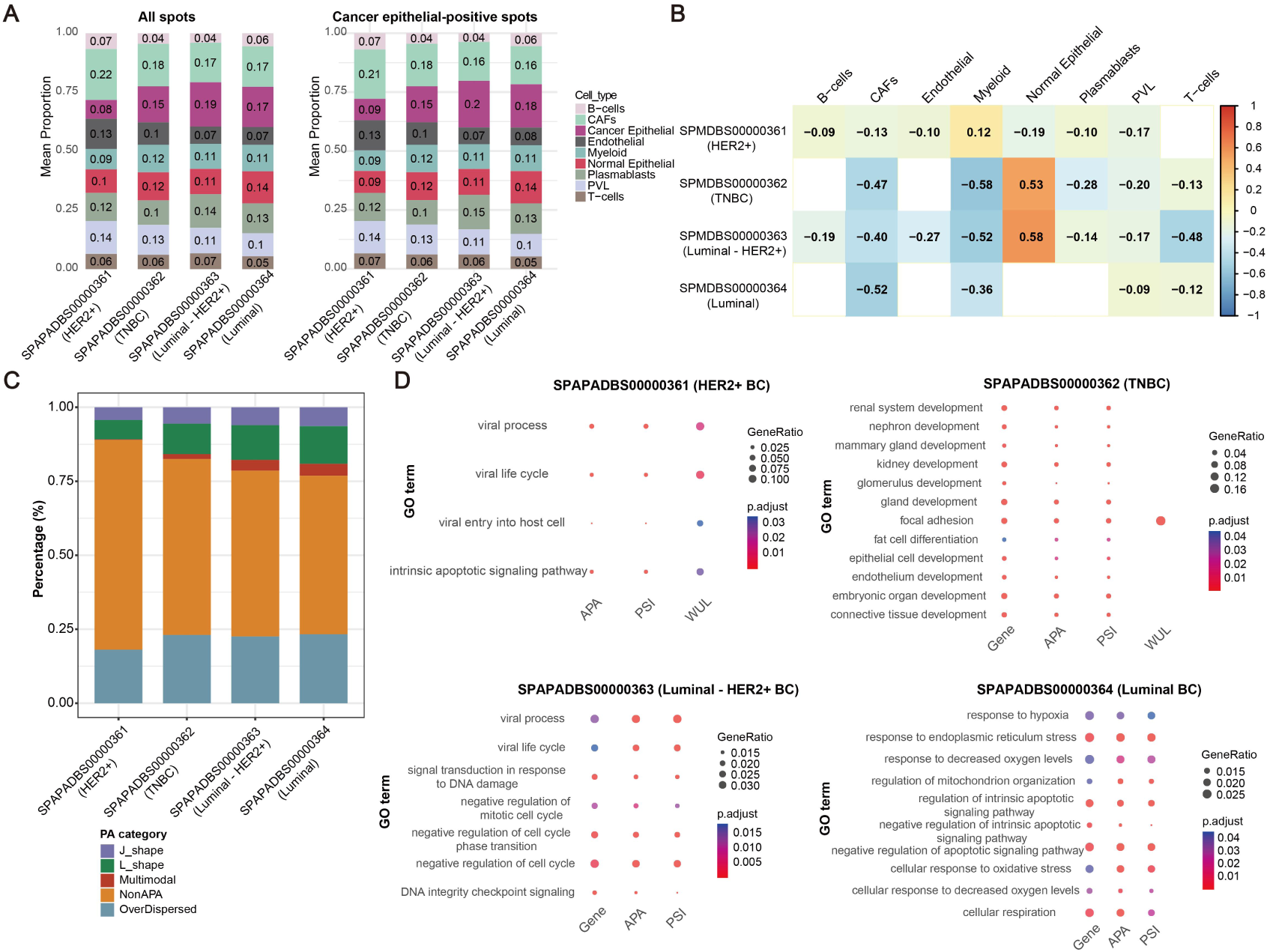
Distinct distributions and functional characteristics of cancer epithelial cells across breast cancer (BC) subtypes. **(A)** Cell composition of all spots and cancer epithelial cell–enriched spots across four major BC subtypes: HER2+ BC (SPAPADBS00000361), triple-negative BC (TNBC, SPAPADBS00000362), Luminal-HER2+ BC (SPAPADBS00000363), and Luminal BC (SPAPADBS00000364). **(B)** Correlation between cancer epithelial cell composition and other cell types in epithelial cell–enriched spots across the four subtypes. **(C)** Distribution of APA categories across the four BC subtypes. **(D)** Gene Ontology (GO) enrichment analysis of gene features, APA events, PSI, and WUL in epithelial cell–enriched spots across the four subtypes.

To investigate the markedly distinct cellular dynamics across different molecular subtypes of breast cancer, further APA analysis revealed subtype–specific APA usage patterns (Figure 5C and Figure S5). For example, HER2+ BC and TNBC show significantly lower levels of the Multimodal APA usage pattern. This suggests that APA may play a crucial role in regulating breast cancer progression. Next, we conducted spatial gene expression and APA analyses, confirming the distinct functional characteristics of cancer epithelial–enriched spots in each BC subtype (Figure 5D). For example, in TNBC, cancer epithelial–enriched spots were enriched in pathways related to cell development and differentiation, whereas HER2+, Luminal – HER2+, and Luminal BCs showed enrichment in pathways related to viral infections, immune responses, oxidative stress, cell cycle and apoptosis (Figure 5D). This suggests that cancer epithelial–enriched spots in TNBC have a higher stemness and differentiation potential compared to those in other subtypes, which is consistent with the consensus that TNBC is the most aggressive subtype of breast cancer [35, 36]. Thus, our findings underscore the functional heterogeneity of cancer epithelial cells in driving subtype–specific tumor progression and highlight the relevance of leveraging APA patterns as potential biomarkers and therapeutic targets in breast cancer, providing a molecular basis for subtype–specific interventions.

Previous studies [37, 38] have shown that *TSPAN8* regulates the malignant progression of breast cancer; however, its expression patterns across different breast cancer subtypes remain unclear. Our spatial transcriptomic analysis revealed that *TSPAN8* is specifically expressed in cancer epithelial cell–enriched spots of TNBC and undergoes subtype–specific APA at the chr12:71125071 locus (**Figure 6A**, 6B, and Figure S6). Subsequent single–cell transcriptomic analysis of TNBC further validated this finding and demonstrated that *TSPAN8* serves as a marker gene for a distinct subpopulation of cancer epithelial cells, effectively distinguishing different epithelial cell subgroups (Figure 6C–F).

**Figure 6.**
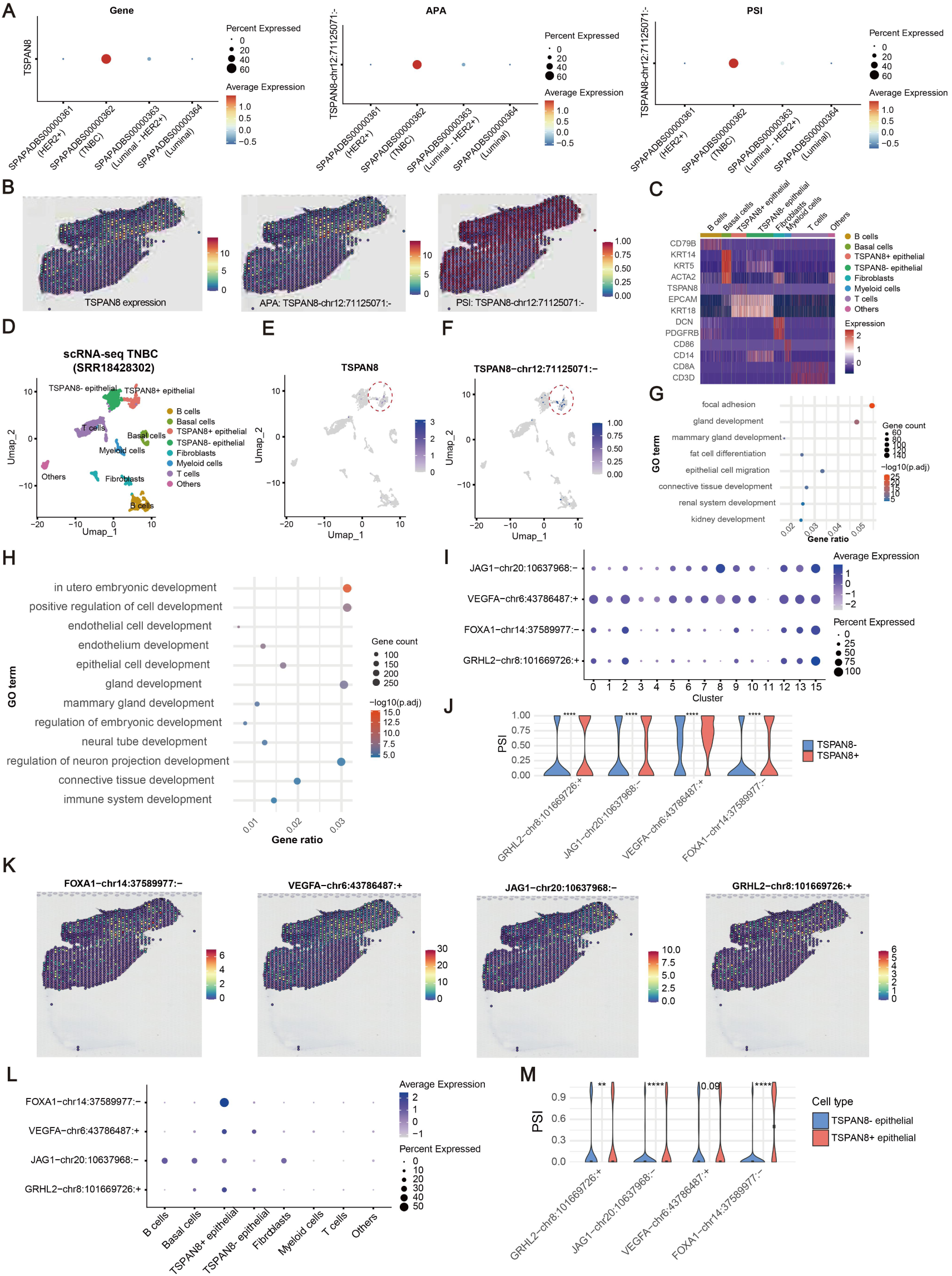
Critical roles of APA in regulating *TSPAN8*-positive cancer epithelial cells in TNBC. **(A)** Dot plots showing expression levels, APA usage, and PSI values of *TSPAN8* in cancer epithelial cell–enriched spots across four breast cancer (BC) subtypes. **(B)** Spatial plots illustrating expression, APA usage, and PSI of *TSPAN8* specifically in TNBC cancer epithelial cell–enriched spots. **(C)** Heatmap displaying cell type–specific marker gene expression in TNBC. **(D)** UMAP plot of single-cell annotations in TNBC. **(E–F)** Feature plots showing *TSPAN8* and *TSPAN8*-chr12:71125071:-expression in a subset of cancer epithelial cells. **(G–H)** GO enrichment analysis of marker genes (G) and genes with dysregulated APA (H) in *TSPAN8*-positive cancer epithelial cells in TNBC. **(I–K)** Dot plots (I), violin plots (J), and spatial plots (K) highlighting APA events of *JAG1*, *GRHL2*, *VEGFA*, and *FOXA1*, which are enriched in the pathways of cell development and differentiation in *TSPAN8*-positive cancer epithelial cells from TNBC spatial transcriptomics. **(L–M)** Single-cell transcriptomic APA analysis in TNBC, with dot plots (L) and violin plots (M) validating the APA events of these genes.

In line with previous studies [37, 38], comparative analyses between *TSPAN8*–positive and *TSPAN8*–negative cancer epithelial cells showed that both differentially expressed genes and APA events were consistently enriched in pathways related to cell development and differentiation (Figure 6G and 6H). Moreover, integrated spatial and single–cell APA analyses in TNBC revealed that genes associated with malignant phenotypes of breast cancer, including *FOXA1*, *VEGFA*, *JAG1*, and *GRHL2*, exhibited markedly distinct expression and APA patterns between *TSPAN8*–positive and *TSPAN8*–negative cancer epithelial cells (Figure 6I–M). Collectively, these results suggest that *TSPAN8*–positive cancer epithelial cells, through unique APA regulation and transcriptional programs, may play a pivotal role in cell development, differentiation, and malignant progression of TNBC.

### A comprehensive online platform for visualizing spatial APA data with gene expression, and cellular dynamics analysis in health and disease

Integrating and analyzing spatial APA, gene expression, and cellular dynamics data can be highly challenging for bench scientists without bioinformatics or programming expertise. To address this, we developed SpatialAPAdb (http://www.biomedical-web.com/spatialAPAdb), a comprehensive online platform designed to simplify these complex analyses (**Figure 7 and Table 1**). With its intuitive search and browsing tools, SpatialAPAdb provides seamless access to a diverse range of relevant multi–omics ST data, empowering researchers to efficiently integrate and analyze spatial APA, gene expression, and cellular dynamics in both healthy and diseased states. Additionally, the platform offers a suite of advanced visualization and analysis tools, enabling users to uncover meaningful insights from these multi–omics data with ease. These features support advances in disease prevention and health maintenance.

**Figure 7.**
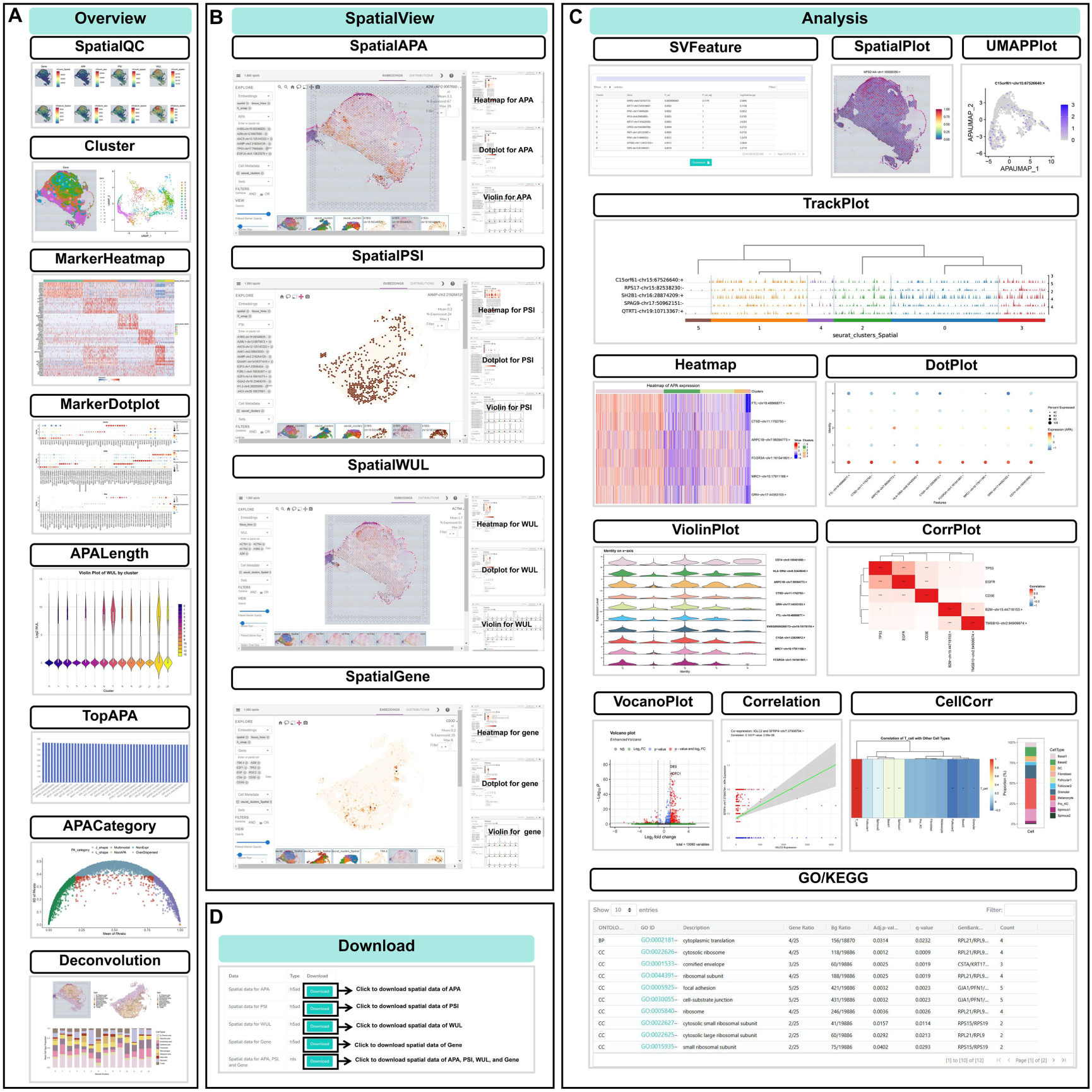
A comprehensive online platform for exploring the spatial APA Atlas. **(A)** The eight visualization tools in the Overview module for assessing data quality and visualizing APA, PSI, WUL, and gene expression features, including their spatial, expression, and length distributions. **(B)** The four interactive tools in the SpatialView module to explore differential expression, diversity, and dynamics of APA, PSI, WUL, and gene expression at spatial resolution. **(C)** The twelve custom analysis tools in the Analysis module to study the spatial diversity and dynamics of APA, PSI, WUL, gene expression, and cell composition, as well as their co-expression relationships and biological functions. **(D)** The Download module for accessing and exporting data.

**Table 1.**
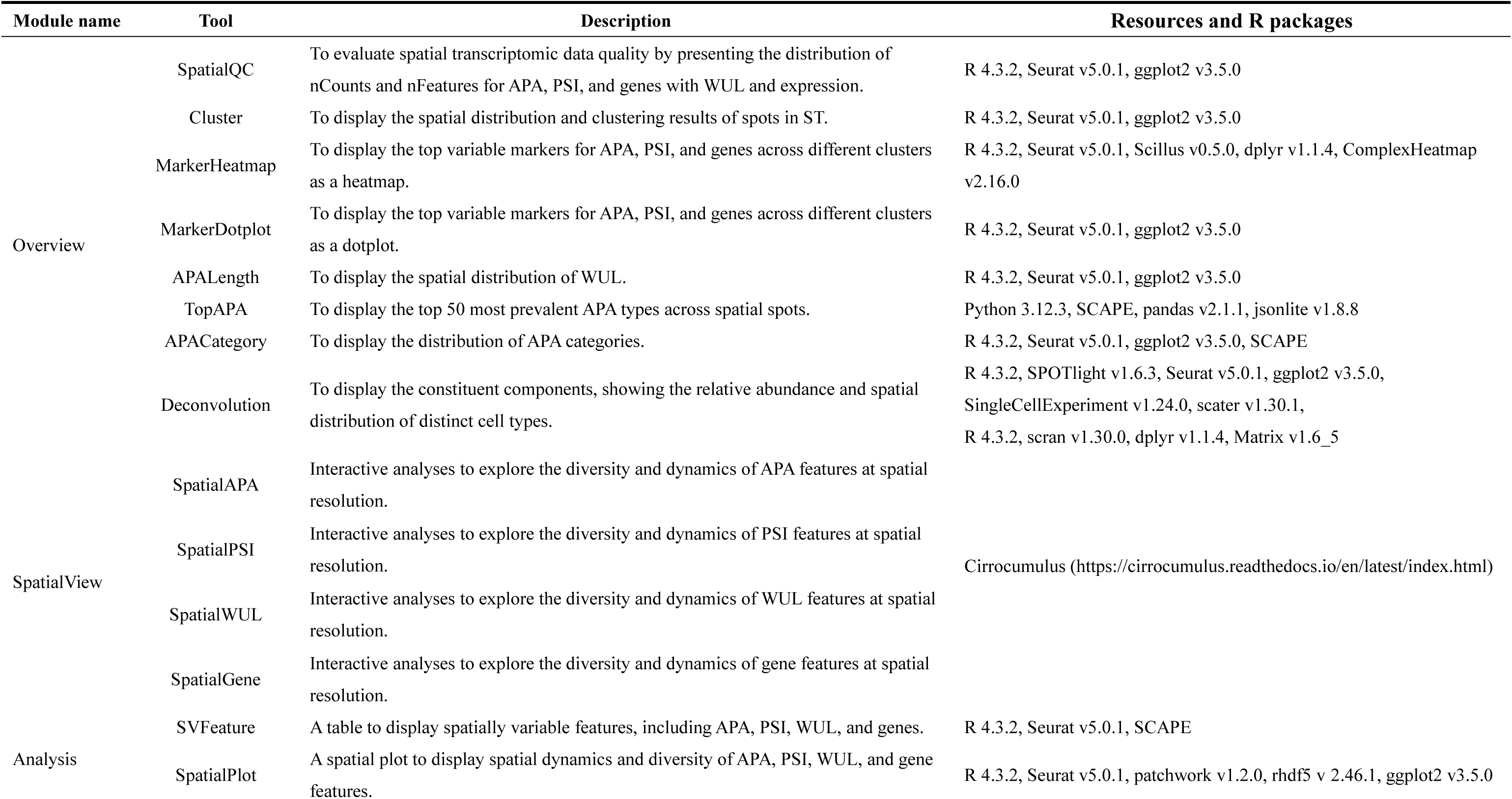
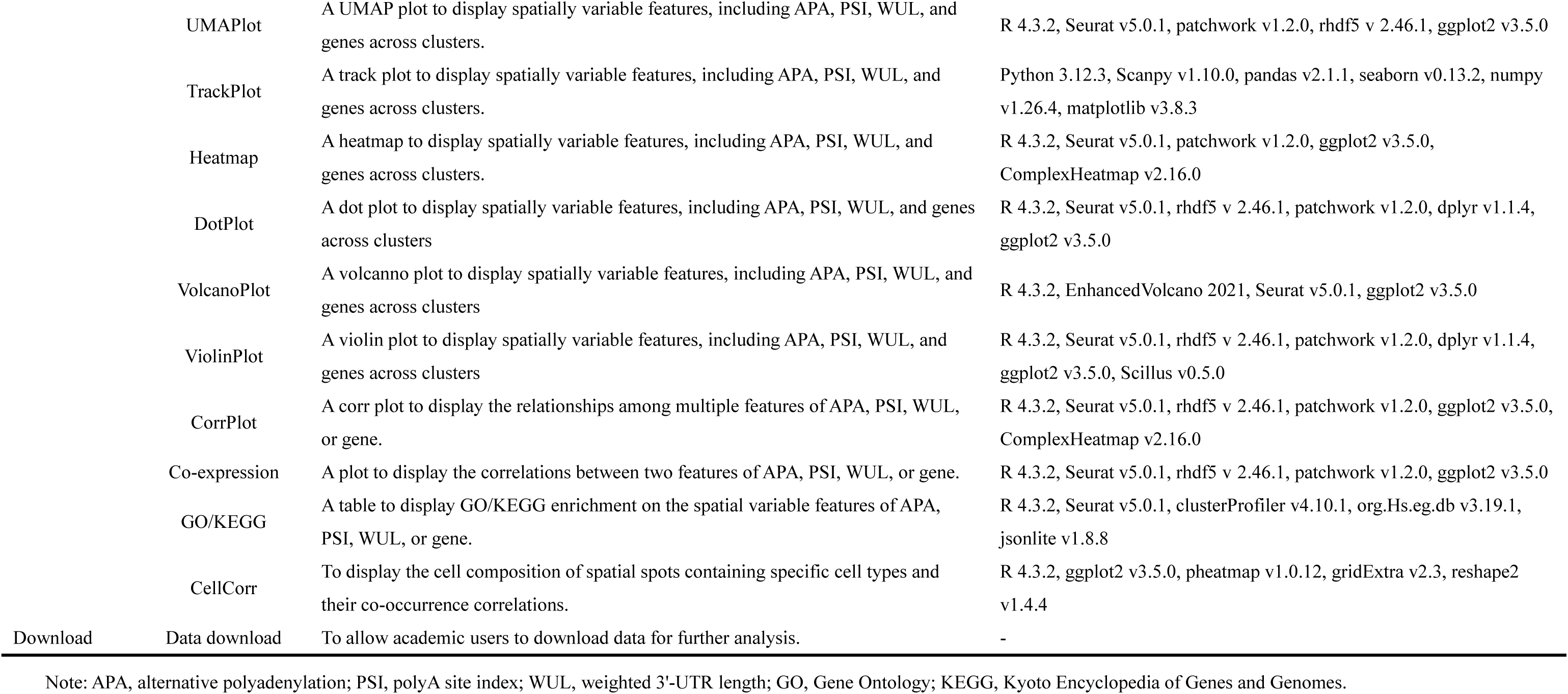
Details of the implementation and functionality of the advanced visualization and analysis tools in SpatialAPAdb.

The platform’s tools are organized into four main modules: Overview, Spatial View, Analysis, and Download (Figure 7 and Table 1). The Overview module includes eight tools for evaluating data quality and visualizing the distribution of APA, PSI, WUL, and gene expression, as well as integrating deconvolution techniques to reveal the spatial distribution of different cell types within tissues (Figure 7A). The Spatial View module provides interactive tools to explore differential expression patterns, as well as the diversity and dynamics of APA, PSI, WUL, and gene expression at a spatial resolution (Figure 7B). The Analysis module contains 12 tools for studying spatial diversity, dynamics, relationships, and biological functions of APA, PSI, WUL, gene expression, and cell composition (Figure 7C). High–quality charts generated by these tools are available for free download, allowing integration into researchers’ publications. Additionally, all data in SpatialAPAdb is publicly accessible for academic use, promoting global scientific collaboration and discovery (Figure 7D).

Comprehensive user guidance and video tutorials are available on the Help page (http://www.biomedical-web.com/spatialAPAdb/help). SpatialAPAdb has been extensively tested across major web browsers, including Google Chrome (version 130.0.6723.117), Safari (version 16.2), Microsoft Edge (version 130.0.2849.80), and Firefox (version 116.0.3), ensuring optimal performance.

With its robust analysis and visualization capabilities, SpatialAPAdb offers a valuable platform for studying the spatial dynamics of APA data, gene expression, and cellular interactions within tissues. This innovative platform not only deepens our understanding of disease mechanisms but also aids in the identification of potential diagnostic and therapeutic targets, highlighting its significant academic and practical value.

## Discussion

This study introduces the SpatialAPA framework, which supports multiple mainstream APA identification methods—including SCAPE v1.0 [17], scAPAtrap v1.0 [20], Infernape v1.0 [31], and scUTRquant v0.5.0 [16]—enabling more flexible and robust detection of spatial APA events across diverse tissue types. Moreover, applying the SpatialAPA framework, we firstly constructed the spatially resolved APA atlas covering 18 human organs and 76 disease states. This atlas provides novel insights into gene expression regulation and its role in health and disease. By integrating spatial cell deconvolution techniques, SpatialAPA identifies the distribution of cell types within tissues, revealing spatially distinct APA regulatory events. The framework uses RPCA to correct batch effects, ensuring consistent clustering and enabling accurate comparisons of APA dynamics across physiological and pathological samples. By integrating ST data, this spatial resolved atlas offers new perspectives on the role of APA in fine–tuning gene regulation, as well as its complex relationship with gene expression and cellular dynamics.

Our findings highlight that APA regulation is highly complex and variable across tissues and disease states. Analysis of 56 publicly available ST projects revealed significant spatial differences in APA, with spatial patterns closely linked to tissue homeostasis and disease progression. We noted that 10x Visium data, while providing valuable spatial context, has inherent resolution and sensitivity limitations, which can affect APA detection in heterogeneous tissues or regions with low cell density. To address this, SpatialAPA applies APA site window correction and spot–level filtering to improve the reliability of APA quantification. On a genome–wide scale, APA events showed stronger organ specificity than gene expression patterns, further supporting the role of APA in the precise regulation of gene expression. Moreover, integrating single–cell RNA–seq data allows spot deconvolution and cell type–specific assignment of APA events, enhancing detection accuracy and enabling more precise interpretation of APA programs in rare or low–density cell populations. The spatially resolved APA atlas provides high–resolution insights into how tissue structures influence gene expression and reflect the molecular mechanisms underlying various diseases.

We used the spatial APA atlas to analyze breast cancer subtypes and found marked heterogeneity in cellular composition and interactions among cancer epithelial cells, CAFs, myeloid cells, and normal epithelial cells. In TNBC, *TSPAN8*–positive epithelial cells showed distinct APA regulation at the chr12:71125071 locus, linked to unique transcriptional programs and enrichment of developmental and differentiation pathways. These cells also exhibited divergent expression and APA patterns of malignancy–associated genes such as *FOXA1*, *VEGFA*, *JAG1*, and *GRHL2*, underscoring their role in TNBC progression and heterogeneity; however, experimental validation is required.

Our findings reveal that APA modulates gene expression and cell–state dynamics in TNBC, promoting stemness, differentiation potential, and functional diversity of cancer epithelial cells. Consistent with prior studies implicating *TSPAN8* in malignancy [37, 38], our results suggest that APA–driven regulation of *TSPAN8*–positive epithelial cells may underlie aggressive tumor behavior and poor prognosis, providing clinical opportunities. Distinct APA signatures and transcriptional programs in these cells could serve as biomarkers for patient stratification, early detection, and disease monitoring, while targeting APA–regulated pathways may open new therapeutic avenues. More broadly, the spatial APA atlas represents a resource for precision oncology. By mapping cell type– and subtype–specific APA events, it can help identify patient–specific vulnerabilities, predict therapeutic response, and guide tailored treatments that integrate genetic and post–transcriptional regulation.

Additionally, we developed the SpatialAPAdb platform, a user–friendly tool for exploring the spatial APA data with gene expression and cell dynamics. This platform makes spatial multi–omics analysis accessible to researchers without bioinformatics expertise. Unlike stAPAminer [15], which lacks batch correction and does not integrate spatial gene expression with cellular dynamics, SpatialAPAdb ensures reliable sample comparisons and offers enhanced analytical capabilities. Compared to existing APA resources (e.g., APADB [6], Animal–APAdb [8], SNP2APA [10], APAatlas [9]), which mainly rely on RNA–seq or 3’–seq data, our study integrates ST data to provide a comprehensive view of APA’s spatial heterogeneity and regulation. Unlike single–cell APA resources (e.g., scAPAatlas [15], scAPAdb [16], scTEA–db [17]), which lack spatial context, SpatialAPAdb preserves tissue architecture and provides high–resolution APA distribution, offering a more refined spatial perspective.

Despite its contributions, several limitations remain. Our analysis relied on ST dataset from specific tissues and disease states, which constrains generalizability; expanding to additional tissues and diseases will broaden its scope. Integration with other high–throughput platforms, such as single–cell RNA sequencing, could further enhance insights into APA regulation. Although the framework is robust, more advanced computational strategies, including deep learning, are needed to improve APA detection accuracy in complex tissues. Potential biases in spatial transcriptomics data and technical constraints of APA detection should also be acknowledged, as they may affect resolution and reliability. Moreover, cross–platform integration remains challenging due to variability in sequencing depth, resolution, and pipelines. Cross–species comparisons may reveal conserved mechanisms and evolutionary trends, offering deeper insights into APA biology. Finally, to address data sparsity, we tested existing imputation methods but found them suboptimal, underscoring the need for tailored solutions. As future work, we aim to develop a deep learning–based imputation framework to improve signal recovery while preserving biological fidelity.

In conclusion, this study introduces the SpatialAPA framework, a robust tool for spatially resolved APA analysis that integrates spatial APA data with gene expression and cellular dynamics. The resulting atlas reveals organ–specific APA patterns, uncovers critical cell–type interactions in diseases like breast cancer, and provides an online platform for exploring tissue– and cell–specific gene regulation. This work enhances our understanding of gene regulation in health and disease and offers new opportunities for therapeutic interventions based on spatial multi–omics data.

## Materials and methods

### Data collection and preprocessing

Despite the availability of various ST platforms, the 10x Visium platform remains the most widely used. To avoid complications arising from data heterogeneity, this study focused exclusively on 10x Visium data. A systematic search of the NCBI SRA database [39] was conducted using the keywords "spatial transcriptomics" and "10x Visium" limited to human data. This initially identified 73 candidate projects. After applying the following inclusion criteria, 56 projects, comprising 363 spatial sections, were remained: (1) exclusion of projects lacking 10x Visium data, (2) exclusion of projects without complete fastq or BAM files, and (3) exclusion of projects without bright–field images.

SRA–format data for these projects were downloaded in batches using the NCBI sratool’s "prefetch" function. For projects failing to download via sratool, the axel tool was used as an alternative. The SRA–format data were converted to fastq format using fasterq–dump, and fastq files for each sample were merged. For projects providing only BAM files, these were manually downloaded and converted to fastq format using Space Ranger’s bam2fastq function to ensure consistency with the reference genome. Bright–field images with spatial information were also downloaded via axel. Images in non–optimal formats (e.g., jpg) or dimensions were processed with Adobe Photoshop CC 2019 to convert to PNG format and resized to at least 2000 pixels. Images with impurities unrecognizable by Space Ranger were manually edited using Loupe Browser 8.

### Implementation of the SpatialAPA framework

The SpatialAPA framework (Figure 1) was implemented within Snakemake [33], leveraging Space Ranger v3.0.0 (https://www.10xgenomics.com/support/software/space-ranger) and integrated multiple single cell APA identification methods, including SCAPE v1.0.0 [17], scUTRquant v0.5.0 [16], scAPAtrap v1.0 [20], and Infernape v1.0 [31]. The Space Ranger was used to process 10x Visium fastq data and brightfield images, utilizing the GRCh38 v44 reference genome from GENCODE [40]. This generated gene expression matrices mapped to spatial coordinates. To ensure data quality, low–quality spots were filtered based on gene count, total read count, and mitochondrial gene proportions, following established criteria [41].

Since each spot on the 10x Visium platform typically contains a mixture of 5–10 cells, directly identifying cell–specific APA events is challenging. To address this, the SpatialAPA framework integrated the SPOTlight v1.6.3 tool [34] for spatial cell deconvolution, enabling the analysis of APA events at the cellular level within their spatial context. A random seed of 42 was used to ensure reproducibility. For reference, high–quality single–cell RNA–seq datasets were curated from the NCBI GEO [42] and HTCA [43] databases. In cases where reference data were unavailable, scRNA–seq datasets from well–established publications were supplemented. Detailed information on the reference datasets used for deconvolution across various tissue types is provided in Supplementary Table S1.

### Construction of a human polyadenylation site reference for benchmark

A comprehensive reference of human polyadenylation (pA) sites was constructed by integrating Gencode v44 [44], PolyASite 3.0 [45], and PolyA_DB [46]. Within each database, pA sites within ±25 nucleotides (nt) were collapsed, retaining the most downstream site per cluster. For cross–database integration, PolyASite 3.0 sites were used as the primary set, followed by addition of non–overlapping (±25 nt) PolyA_DB sites and subsequently Gencode v44 sites absent from the merged set, yielding 18,103,429 non–redundant pA sites. Each site was mapped to terminal exon(s) of Gencode v44 gene models according to genomic coordinate and strand; sites overlapping multiple terminal exons were duplicated to reflect all possible gene assignments. This reference provides a unified resource for alternative polyadenylation analyses in human transcriptomes.

### Benchmark of different APA identification methods in the SpatialAPA framework

Four APA identification tools—SCAPE v1.0 [17], scUTRquant v0.5.0 [16], scAPAtrap v1.0 [20], and Infernape v1.0 [31]—were benchmarked on spatial transcriptomics datasets from five human tissues (Brain, Oral Cavity, Ovary, Pancreas, and Stomach) using default parameters in the SpatialAPA framework. APA sites within each tissue were merged if located within ±25 nucleotides to generate non–redundant sets; for scAPAtrap and Infernape, interval endpoints were used as representative sites. Biological accuracy was assessed by comparing predicted APA sites with the reference set integrating Gencode v44 [44], PolyASite 3.0 [45], and PolyA_DB [46], using bedtools window (±25 nt). Match and non–match rates were calculated as the proportions of overlapping and tool–specific sites, respectively. Computational performance was evaluated using a unified Snakemake workflow, recording runtime and peak memory with execution standardized to 8 threads per tool.

### Spatial APA identification across different human tissues

Compared with other single–cell APA identification methods, SCAPE v1.0 demonstrates higher sensitivity and specificity within the SpatialAPA framework. Accordingly, SCAPE was used to identify spatial APA sites in human tissues. First, the Space Ranger processed the data to generate BAM files, which were then analyzed seting the four single cell methods to identify APA events. 3’–UTR coordinates from the GRCh38 v44 GTF file were extracted and saved as a BED file. Moreover, the apamix function of SCAPE identified and quantified APA sites, generating an APA expression matrix. APA sites were annotated to their corresponding genes and positions using the SCAPE’s *AnnotationSite* function. Only APA sites present in at least 25 spots were retained. The Polyadenylation Site Index (PSI) for each APA site was calculated using the psi function from SCAPE, and APA sites were classified into seven types based on their usage patterns: "Multimodal", "NonExpr", "NonAPA", "L_shape", "J_shape", "OverDispersed", and "UnderDispersed" [47, 48]. To evaluate the impact of APA on gene 3’ UTR length, the Weighted 3’ UTR Length (WUL) score was calculated using a published formula [23]. The resulting APA, PSI, and WUL matrices were integrated with the gene expression matrix for each sample using Seurat v5.0.1, followed by additional filtering and saving as RDS files. Finally, the sceasy::convertFormat function was used to convert RDS files to h5ad format for analysis within the SpatialView module of the database.

### Data integration and batch effect correction

ST data often involves clustering of individual spatial sections, which can complicate the integration of multiple sections. To address this issue, we applied RPCA in Seurat v5.0.1 for organ–level data integration. Datasets from the same tissue organ were first merged, and relevant features were identified using the Seurat’s SelectIntegrationFeatures function. The FindIntegrationAnchors function with RPCA was used for dimensionality reduction, followed by integration using the IntegrateData function with a k.weight parameter set to 50. This approach allowed for successful integration of data from 18 different tissue organs. After integration, the unified dataset was split back into individual sections, and cluster information was transferred to each section for subsequent comparison analysis. After integration, cluster information was transferred back to each section for downstream differential analysis. RPCA was validated by visualizing UMAP plots, performing differential expression analysis, and confirming the biological consistency of the integrated dataset.

### Differential analysis

For differential gene expression, we employed Seurat’s FindAllMarkers() function with a log fold change threshold of 0.25 and a minimum expression in 10% of cells. This approach enabled the identification of genes exhibiting significant differences between cell clusters, which were subsequently visualized using VlnPlot and SpatialFeaturePlot. For differential APA event analysis, we applied the same FindAllMarkers() approach to the apa/apapsi/WUL assay to evaluate cluster–specific APA events using identical thresholds. Significant APA events were further validated through dimensionality reduction methods, including PCA and UMAP, to ensure robust cluster–level resolution and interpretability.

### Implementation of SpatialAPAdb

SpatialAPAdb was implemented with a decoupled front–end and back–end framework, as detailed in our previous studies [49, 50]. The application’s operational and data processing functionalities were implemented using Java, while the analytical environment was configured with R 4.3.2 and Python 3.12.3 using Conda (version 23.10.0). This configuration enhanced the analytical capabilities of SpatialAPAdb. The SpatialView module was developed using the Cirrocumulus plugin (https://cirrocumulus.readthedocs.io/en/latest/index.html). The implementation of the visualization and analysis tools in SpatialAPAdb were detailed in Table 1.Finally, SpatialAPAdb is deployed on an Apache Tomcat server.

## Data availability

SpatialAPAdb is publicly accessible at http://www.biomedical-web.com/spatialAPAdb/ without any login or registration requirements.

## Code availability

The SpatialAPA analysis framework is available at https://ngdc.cncb.ac.cn/biocode/tool/7716 and https://github.com/Omicslab-Zhang/spatialAPA.

## CRediT author statement

**Zehang Jiang**: Conceptualization, Methodology, Validation, Software, Investigation, Data curation, Formal analysis, Writing – original draft, Visualization. **Zhuochao Min**: Methodology, Software, Visualization. Zhanying Wu: Methodology, Investigation, Data curation. **Yubin Chen**: Data curation, Formal analysis. **Huashu Wen**: Data curation, Formal analysis. **Cheng Wu**: Methodology. **Jia Guo**: Investigation. **Ke Si**: Data curation. **Guoying Wang**: Resources. **Shuai Mao**: Data curation. **Weizhong Li**: Supervision, Funding acquisition, and Writing – review & editing. **Binghui Zeng**: Supervision, Funding acquisition, and Writing – review & editing. **Wenliang Zhang**: Conceptualization, Methodology, Resources, Supervision, Funding acquisition, Writing – original draft, and Writing – review & editing.

## Acknowledgments

This study was supported by the National Natural Science Foundation of China (Grant No. 32100513), the National Key R&D Program of China (Grant No. 2021YFF1200903), the Basic and Applied Basic Research Foundation of Guangzhou, China (Grant No. SL2023A04J00291), the Science and Technology Program of Guangzhou, China (Grant No. 2024A04J3341), the Guangdong–Hong Kong–Macau Joint Laboratory for Cell Fate Regulation and Diseases, China (Grant No. 2022LSYS008), and the Guangdong Basic and Applied Basic Research Foundation, China (Grant No. 2021A1515111223 and 2022B1515120077).

## Supplementary material

**Figure S1.**
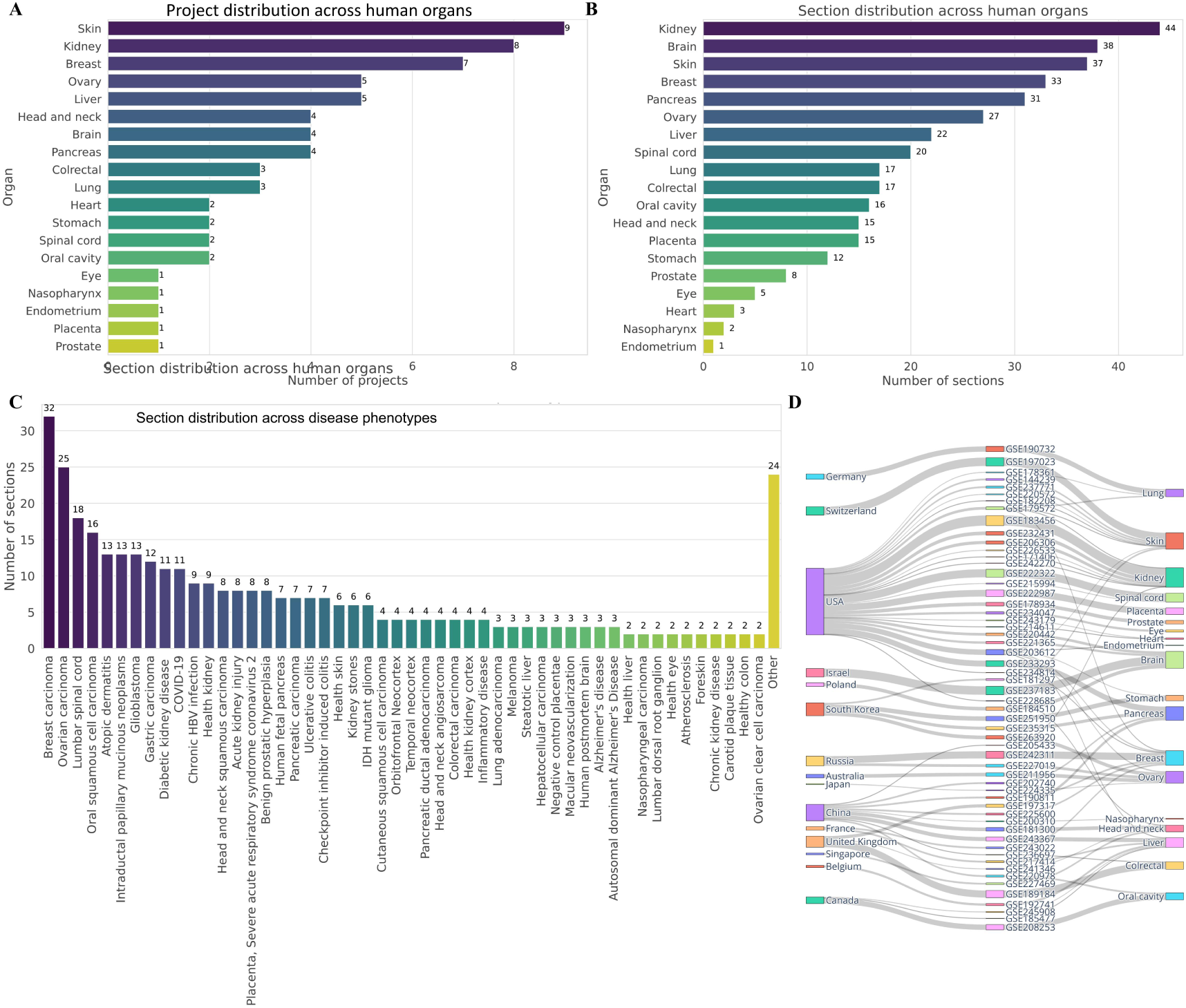
Distribution of spatial transcriptomics (ST) data. **(A)** Distribution of ST projects across various organ types, highlighting those with the most projects**. (B)** Distribution of spatial sections across different organ types. **(C)** Distribution of spatial sections across various disease phenotypes. **(D)** Breakdown of countries and regions contributing to ST data collection, with emphasis on major contributors.

**Figure S2.**
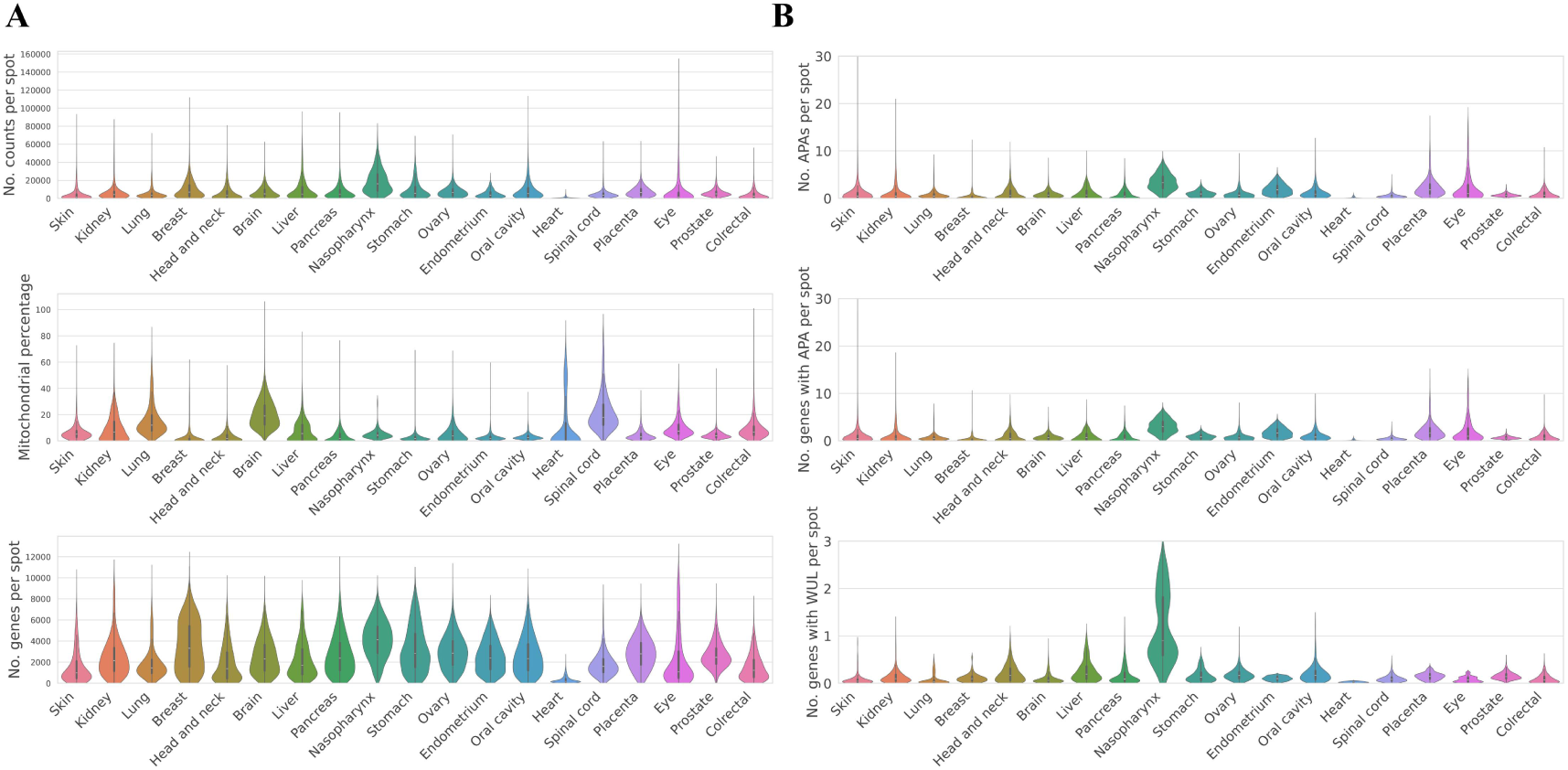
Distribution of genes and APA events per spatial spot across organs. **(A)** Number of genes with APAs per spot in different organs. **(B)** Number of APAs per spot in different organs. **(C)** Number of genes with weighted 3’ untranslated regions (WUL) per spot in different organs.

**Figure S3.**
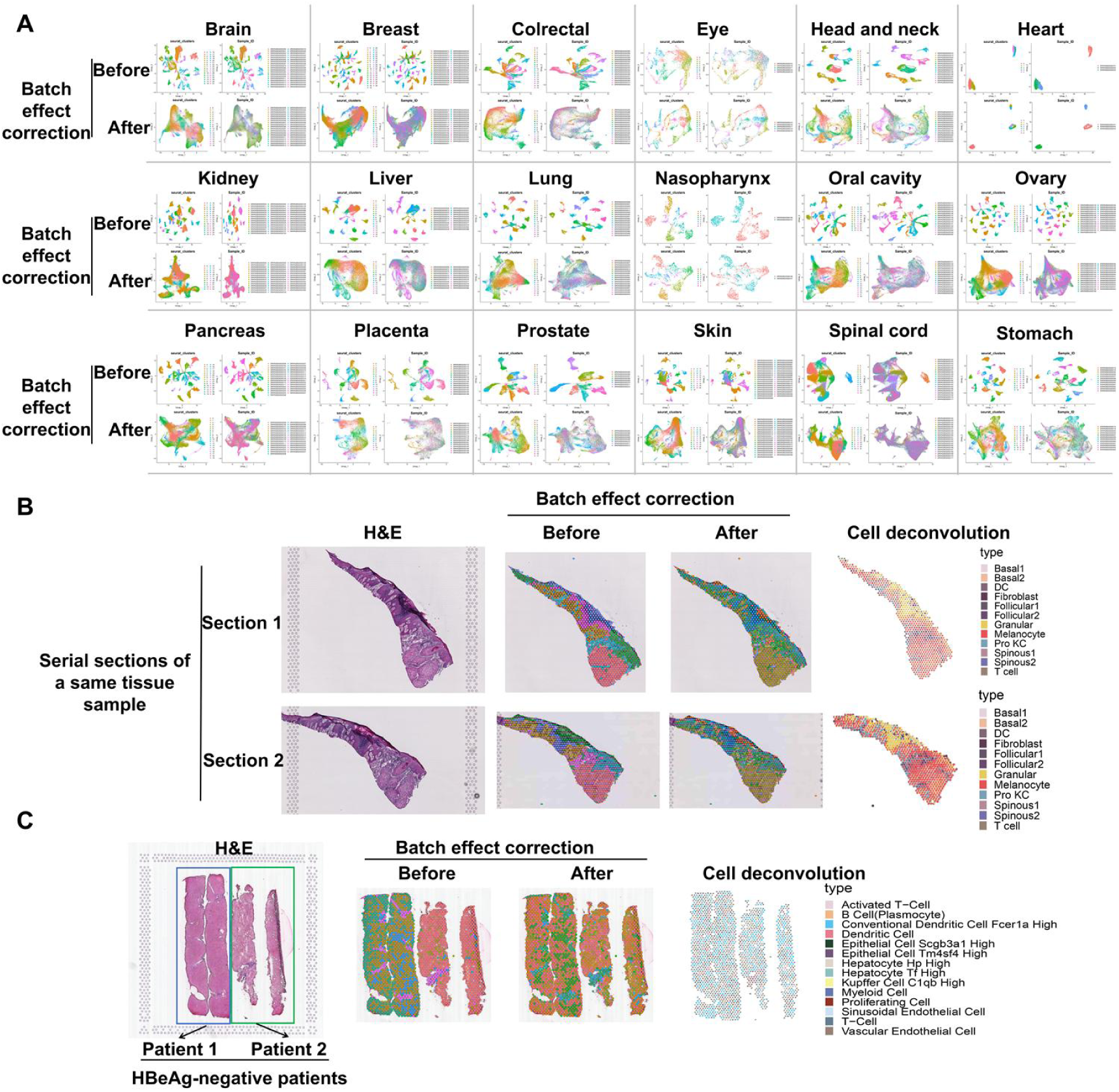
RPCA-based data integration and batch effect correction significantly improve data consistency and enable accurate comparisons across different ST samples. **(A)** Comparison of spatial data integration at the organ level before and after batch effect correction. **(B)** Comparison of spatial spot distribution and cell deconvolution results in two consecutive sections from the same tissue sample before and after batch effect correction. **(C)** Comparison of spatial spot distribution and cell deconvolution results in regions from different tissue samples within the same spatial section before and after batch effect correction.

**Figure S4.**
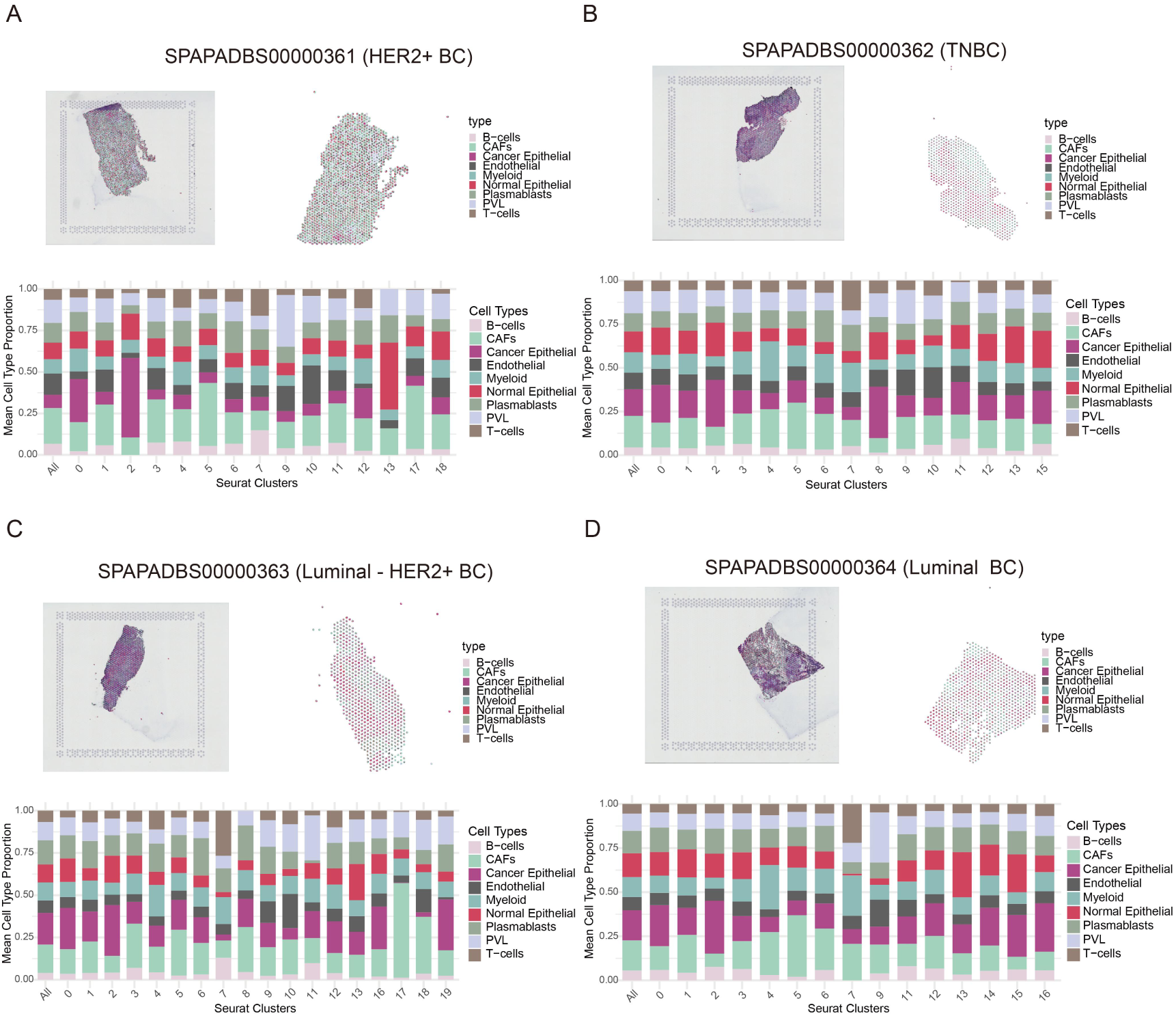
Cell composition across breast cancer subtypes, including *HER2*+ BC (A), TNBC (B), Luminal-*HER2*+ BC (C), and Luminal BC (D).

**Figure S5.**
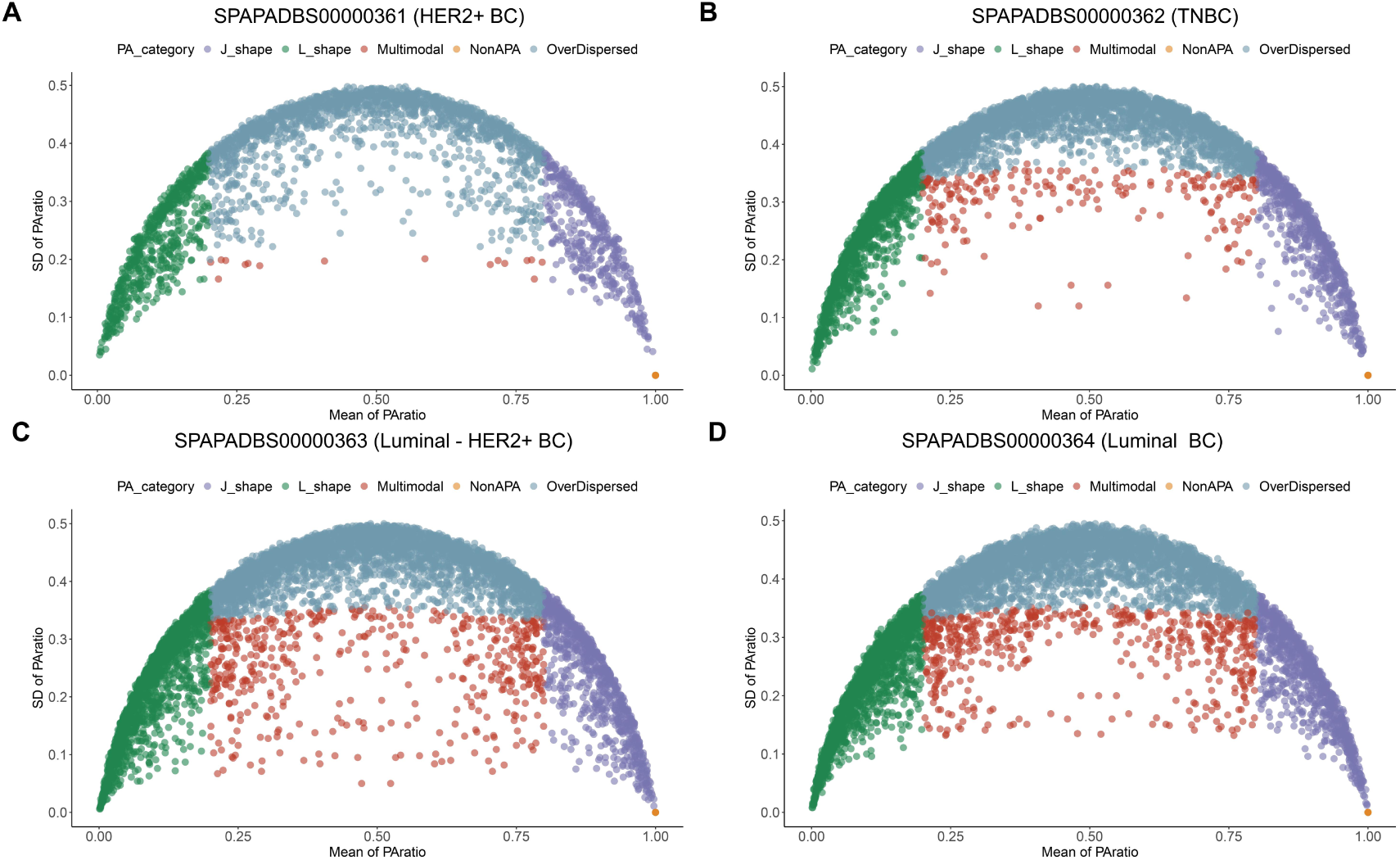
Distribution of APA categories across breast cancer subtypes, including *HER2*+ BC (A), TNBC (B), Luminal-*HER2*+ BC (C), and Luminal BC (D).

**Figure S6.**
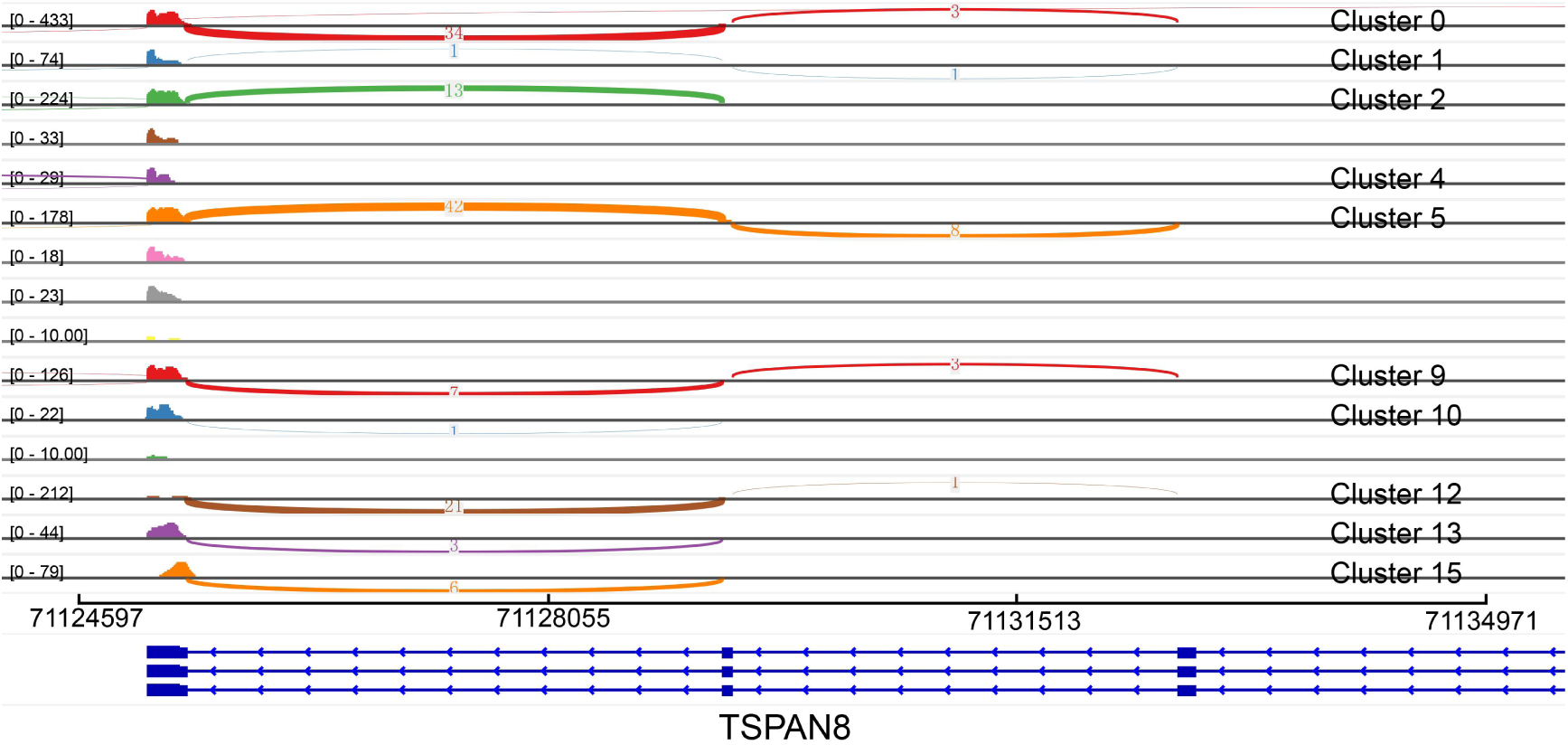
Coverage plots of *TSPAN8*-chr12:71125071:-illustrating cluster-specific alterations of APA events in TNBC.

**Table S1.**
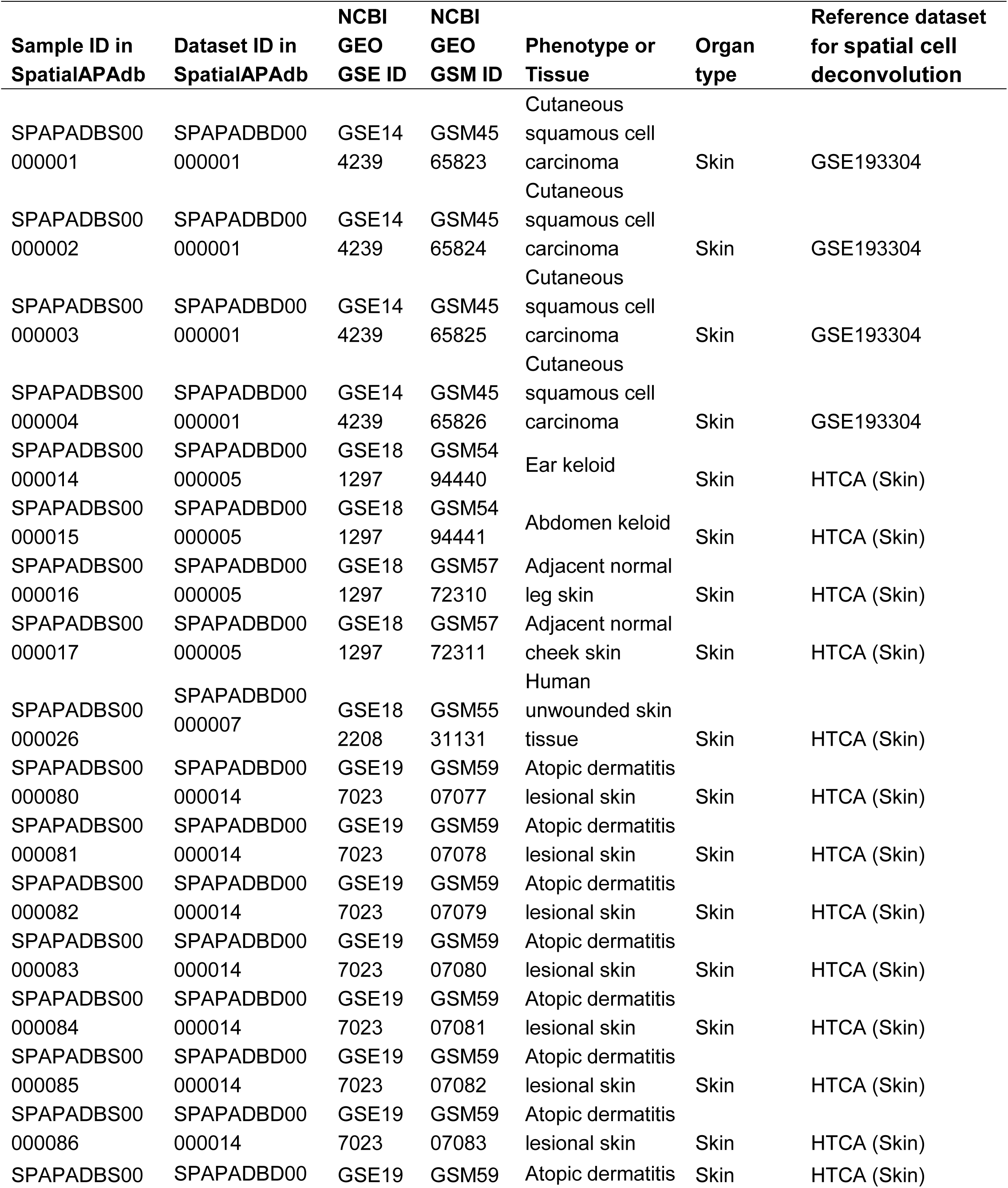
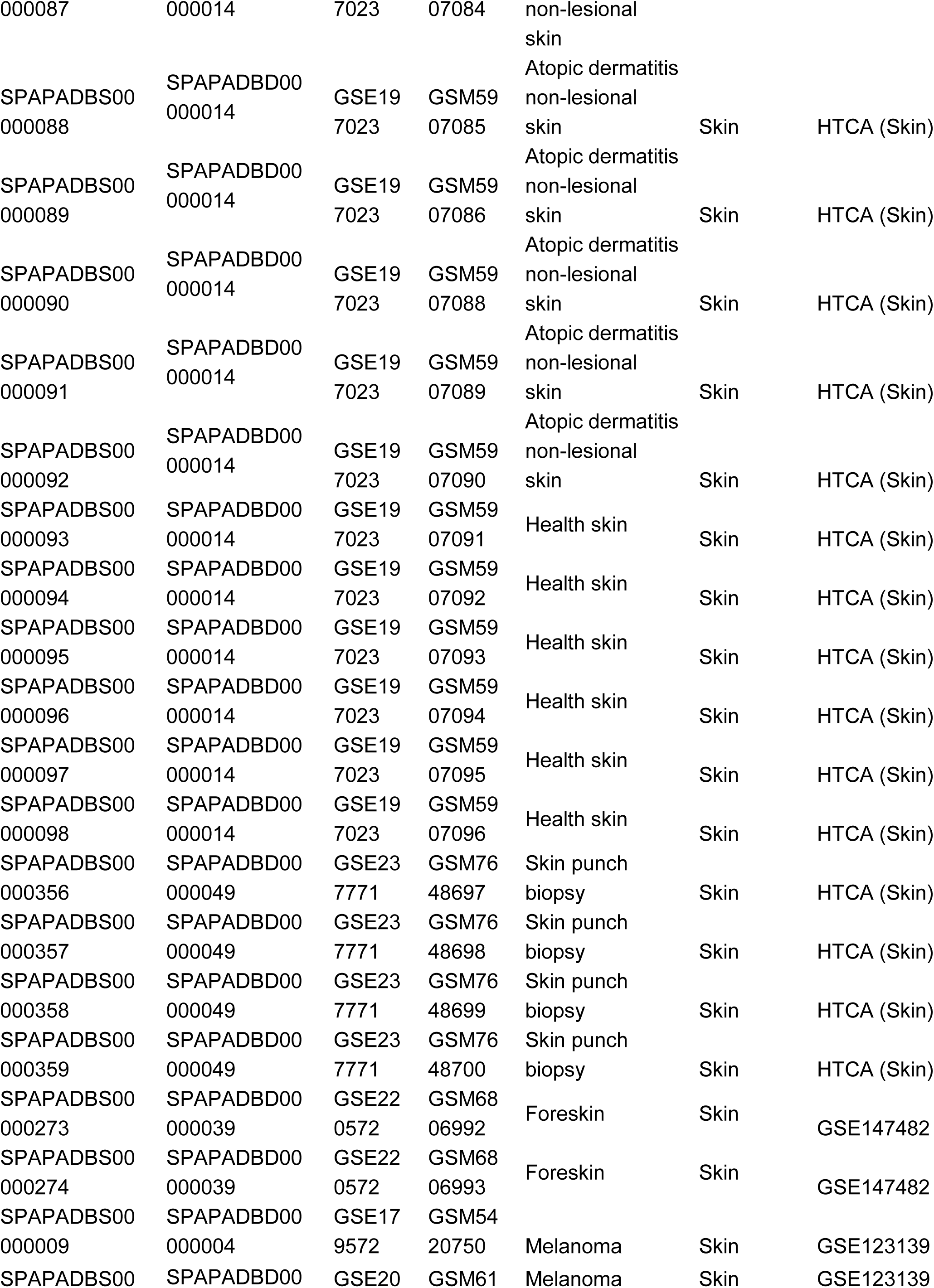
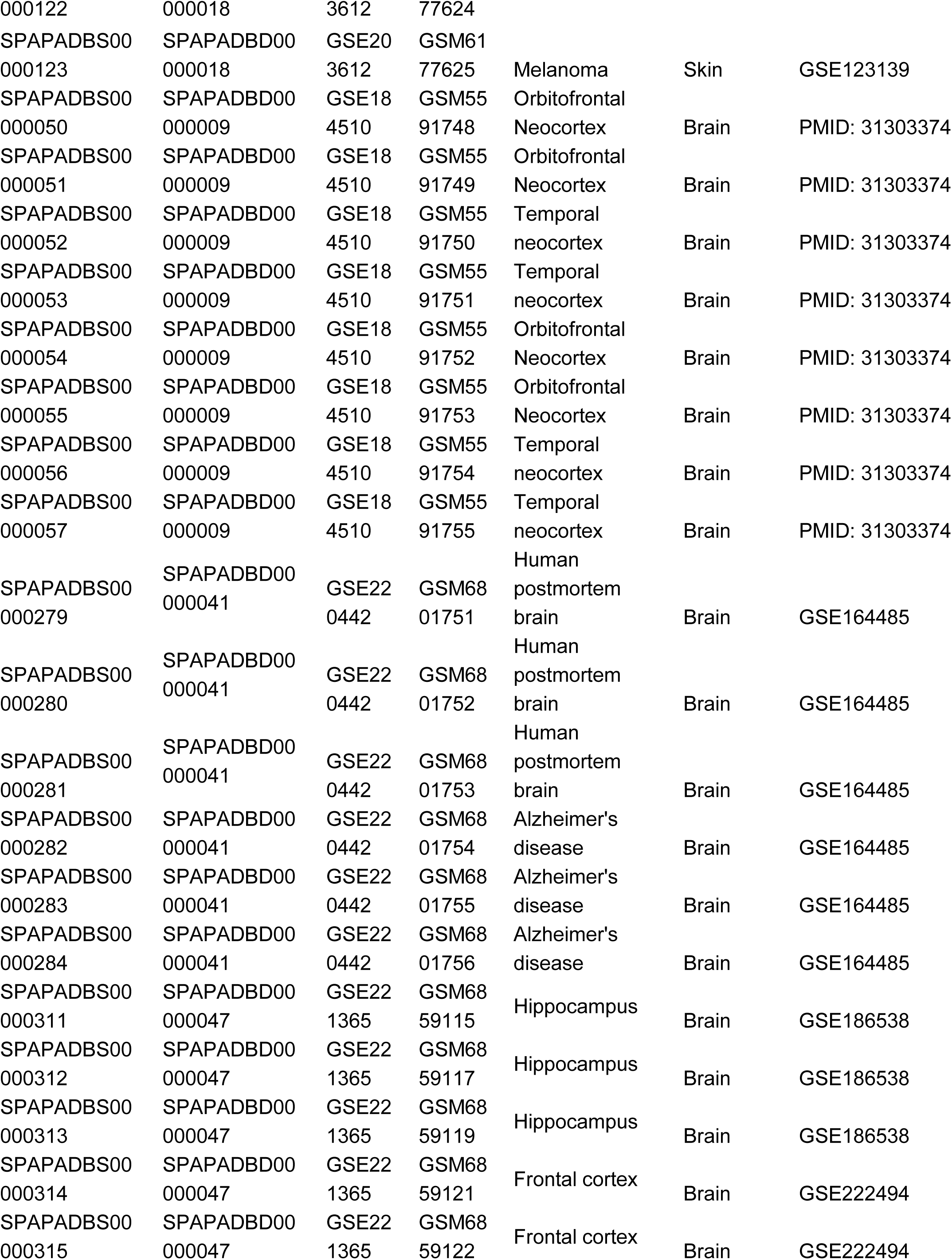
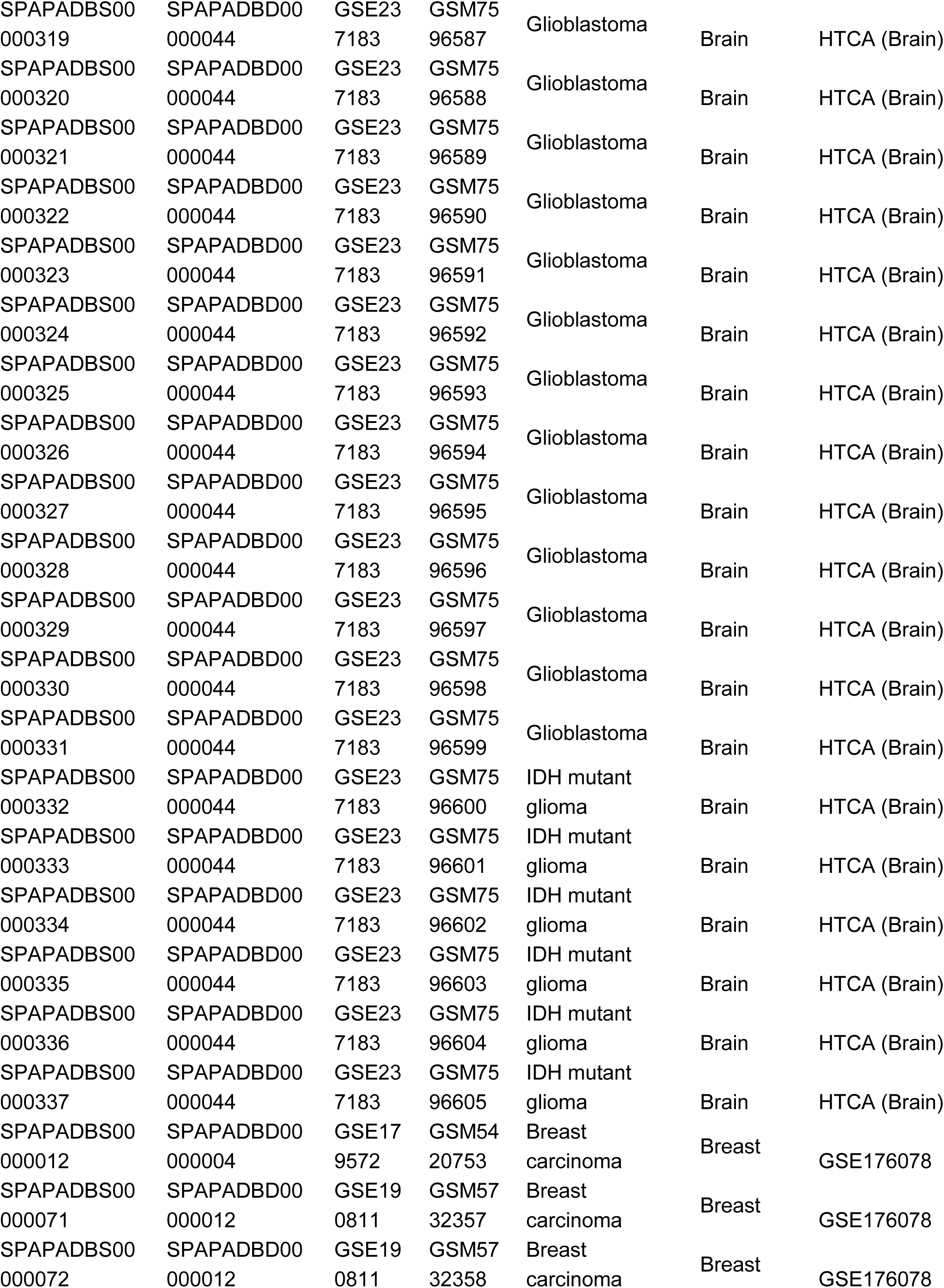
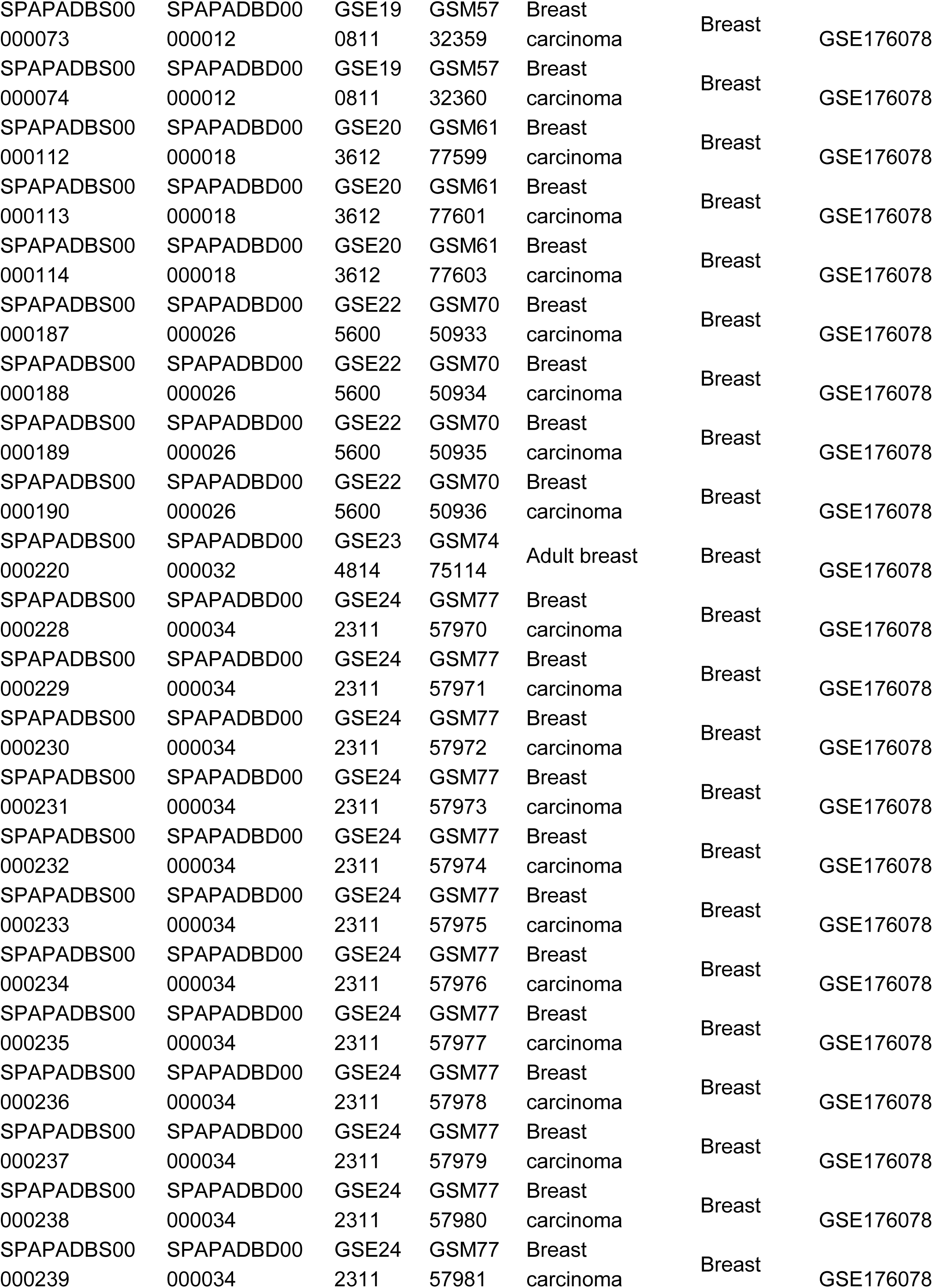
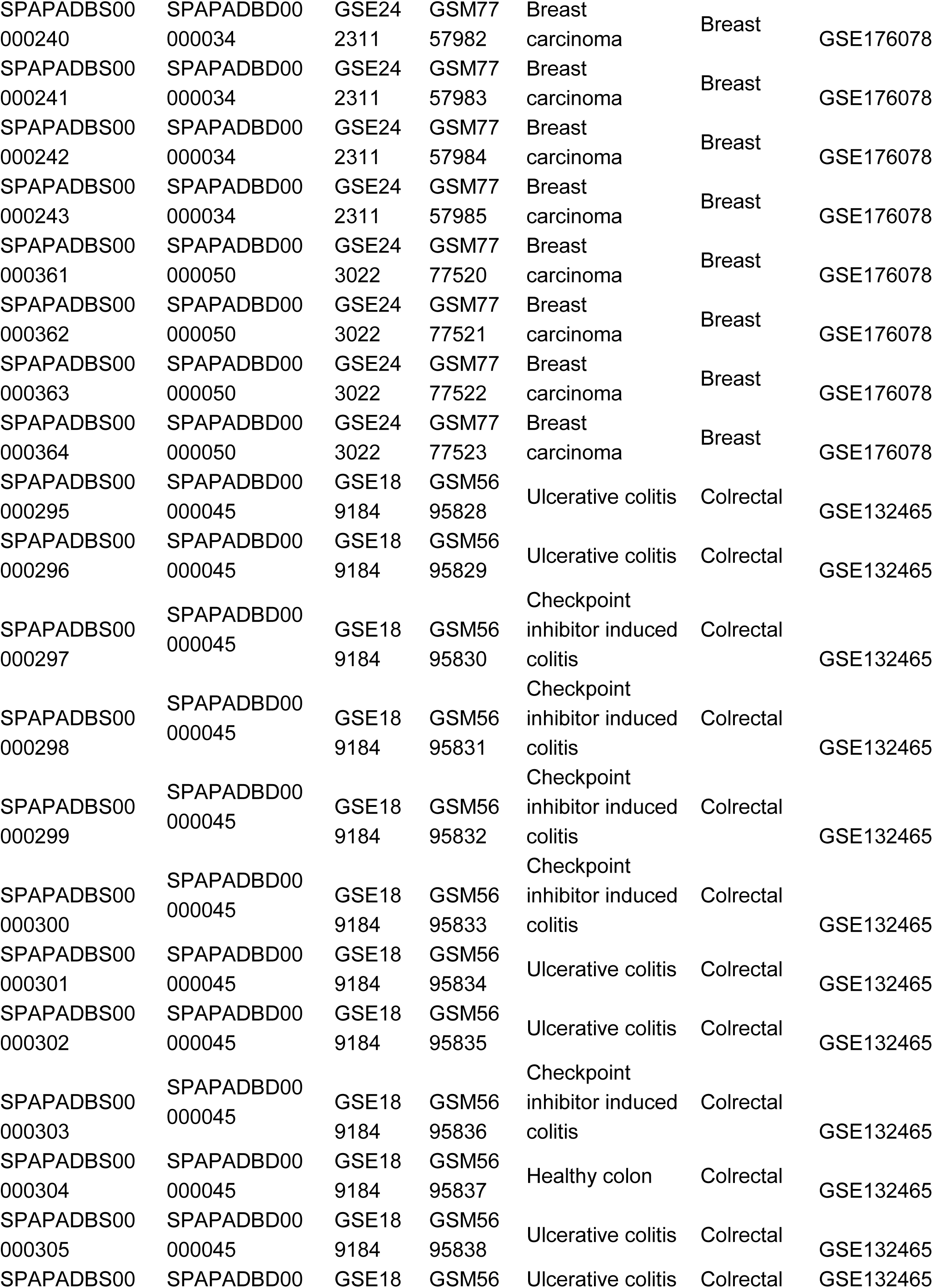
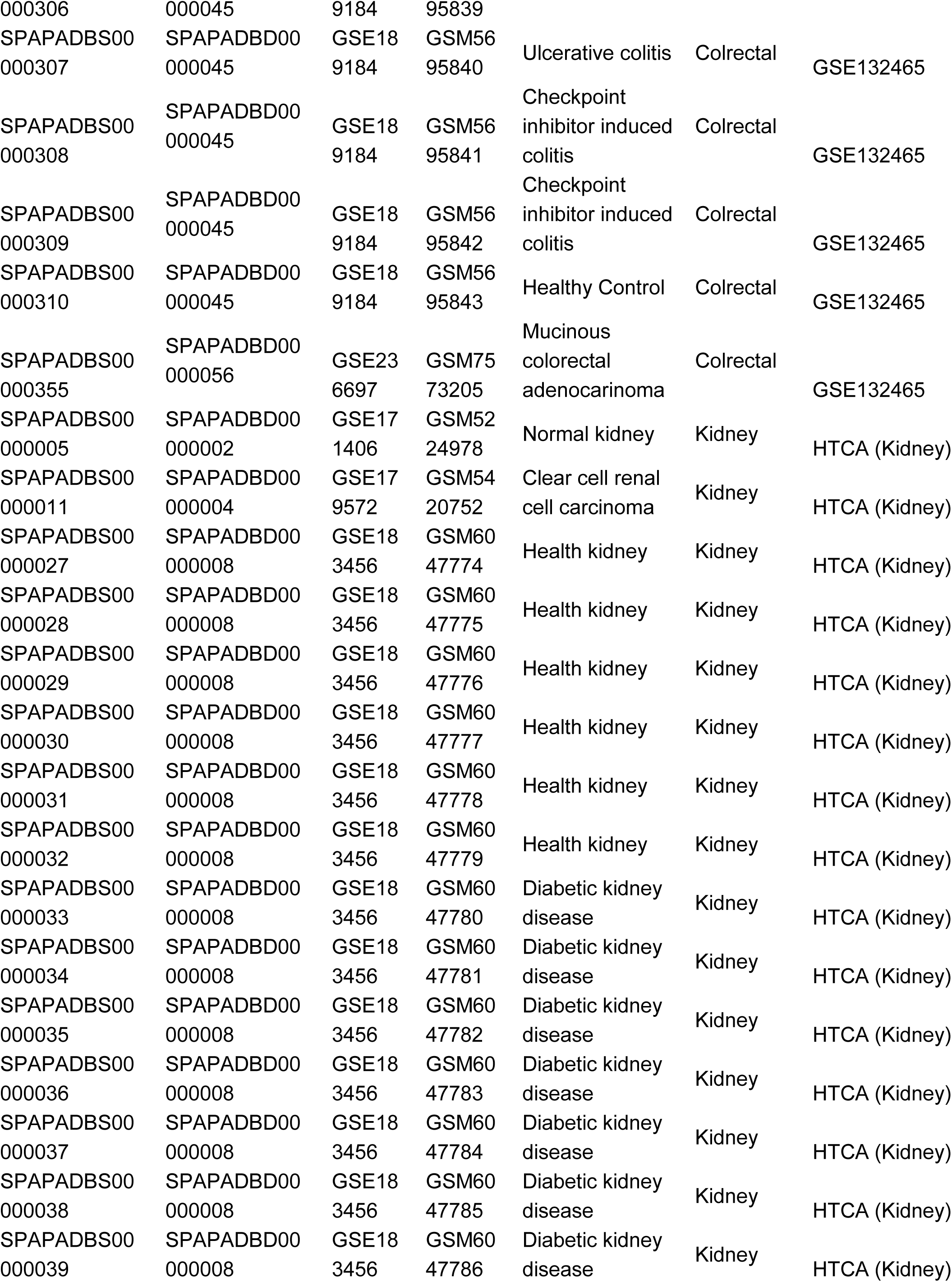
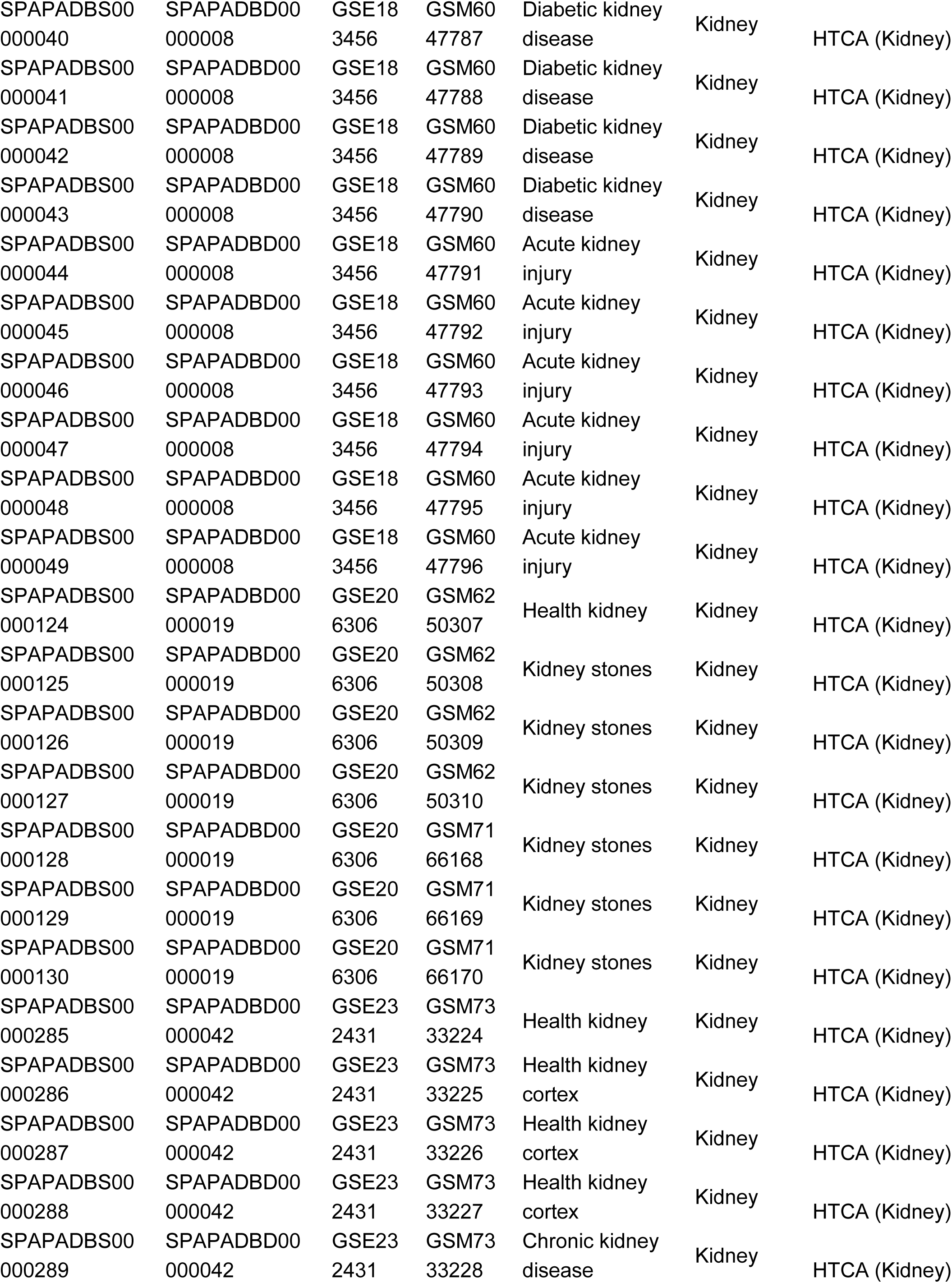
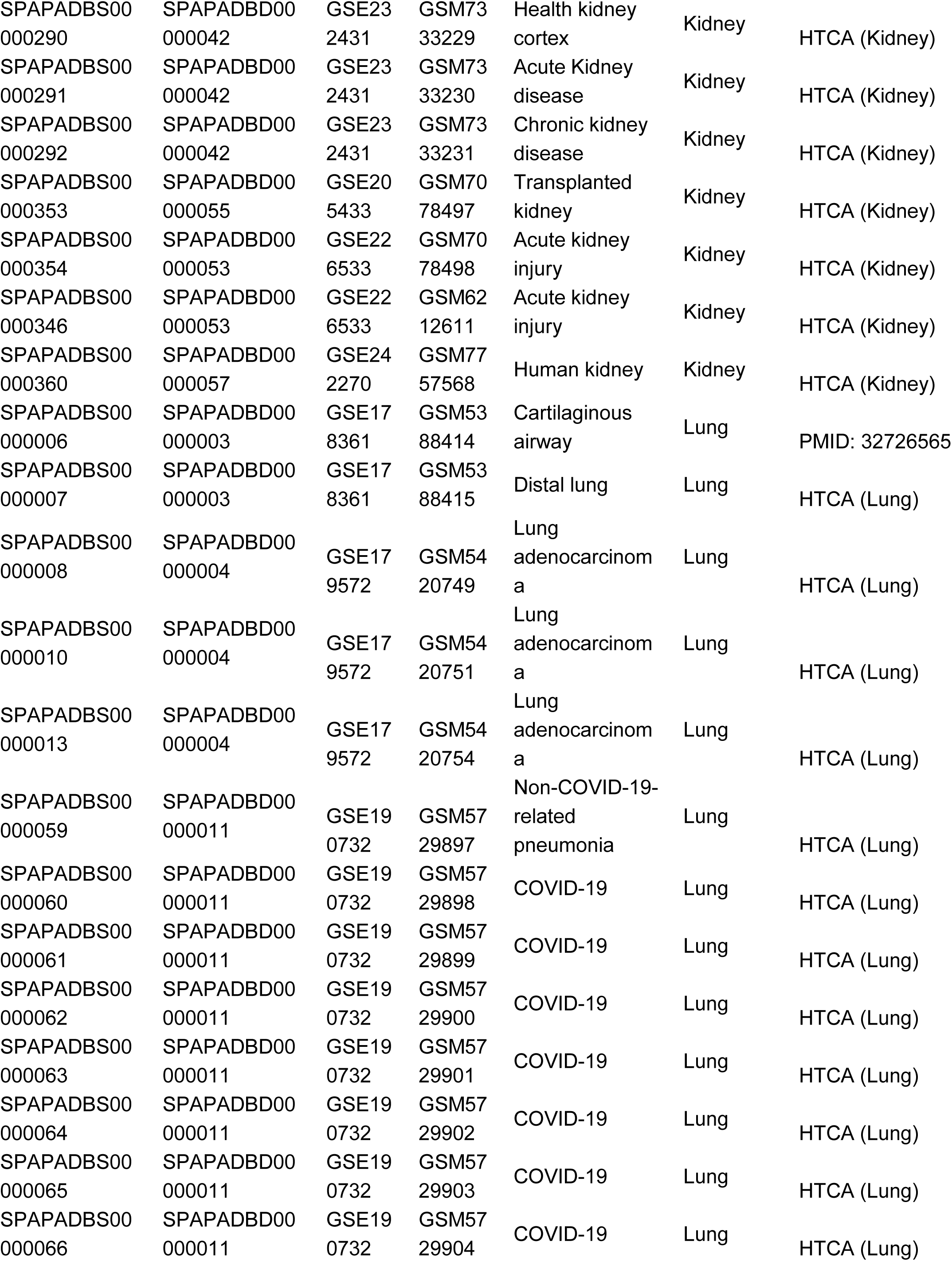
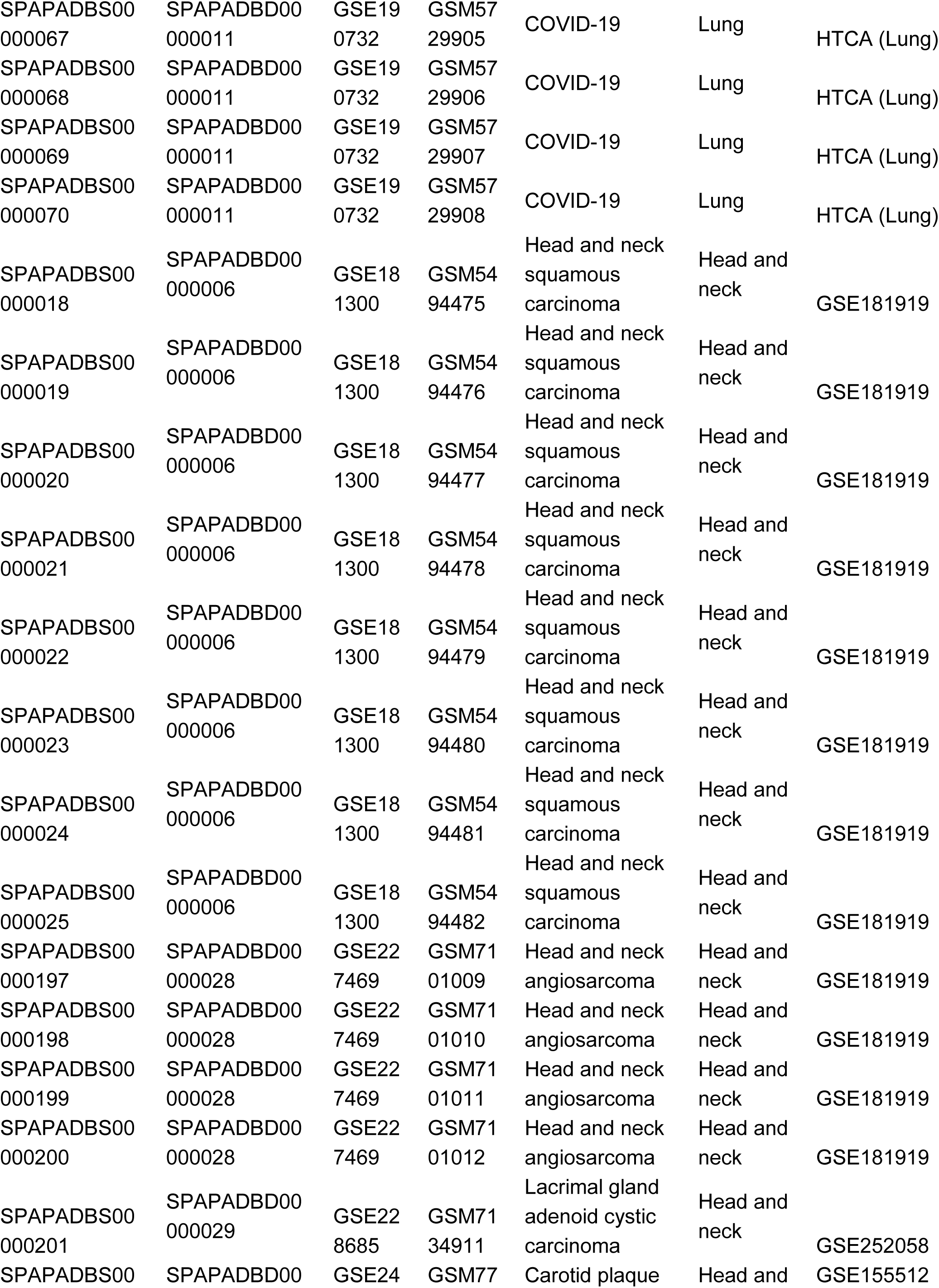
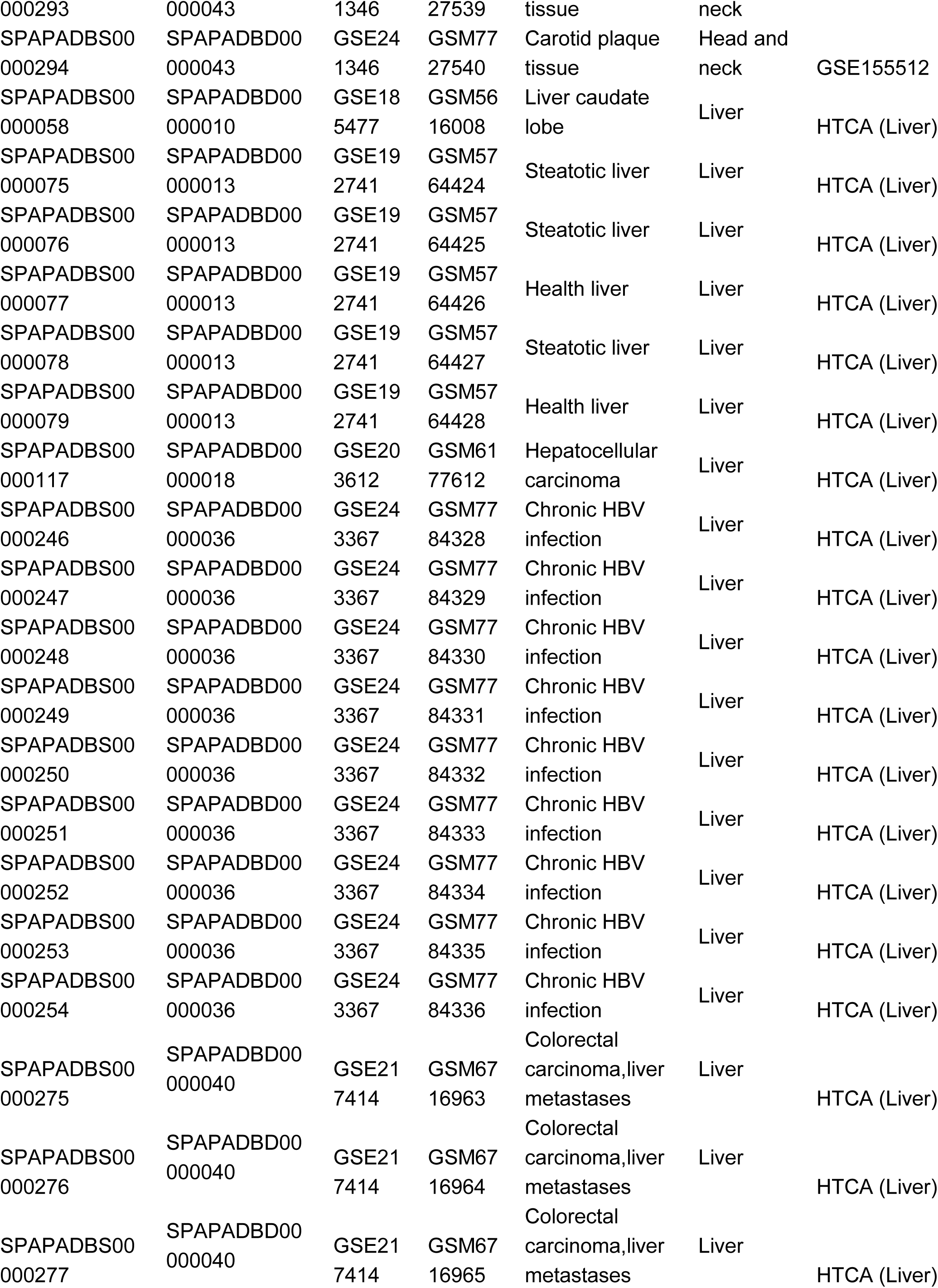
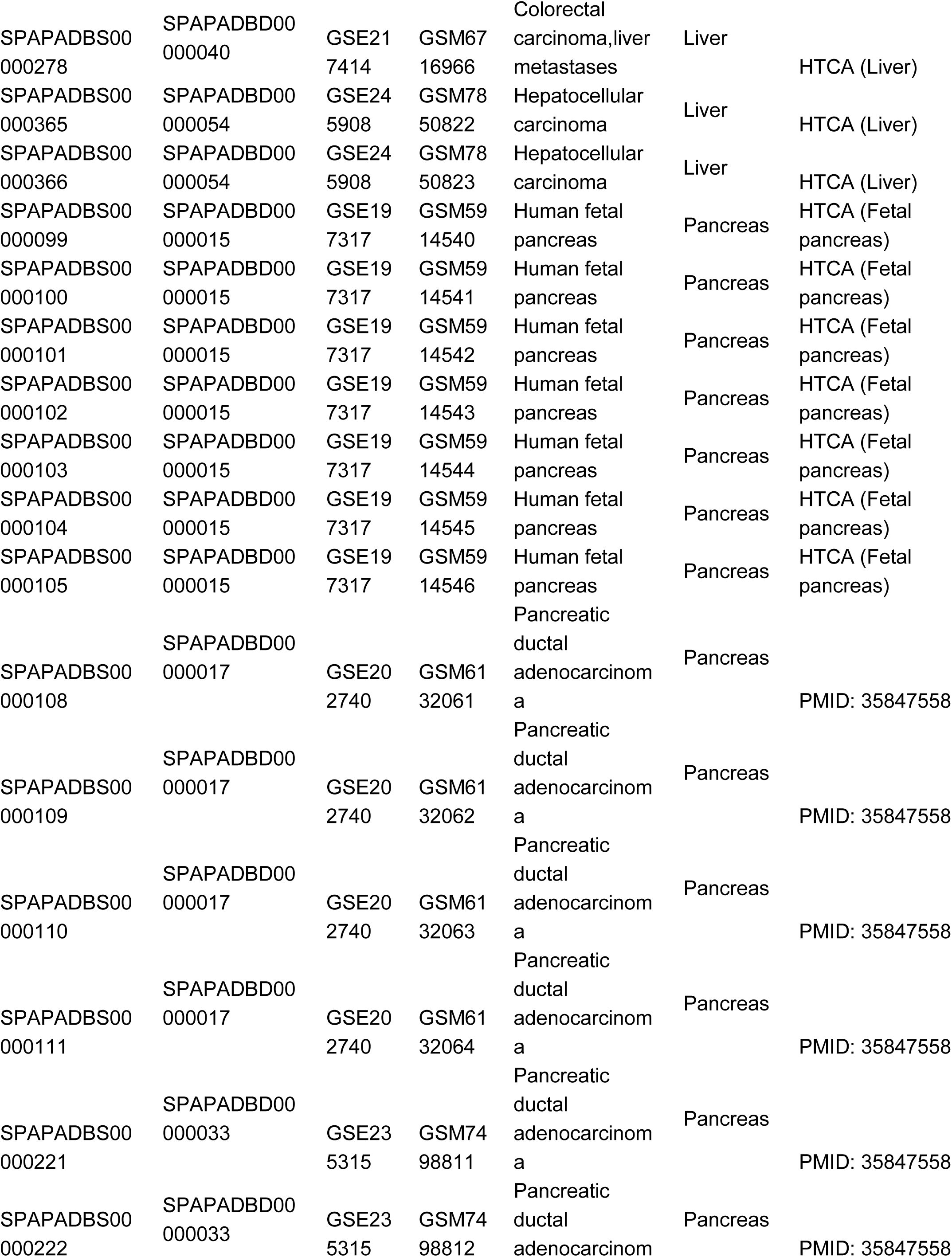
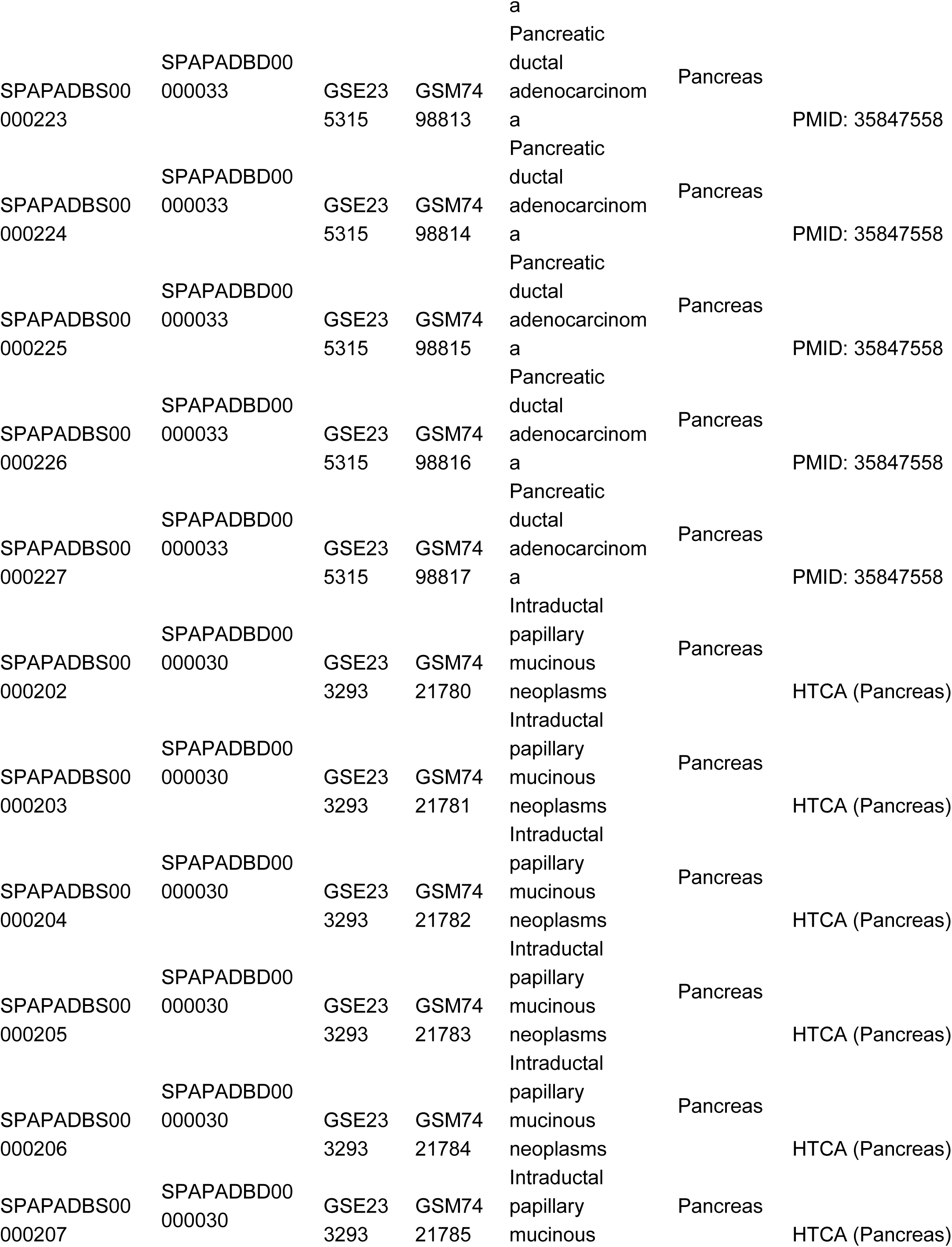
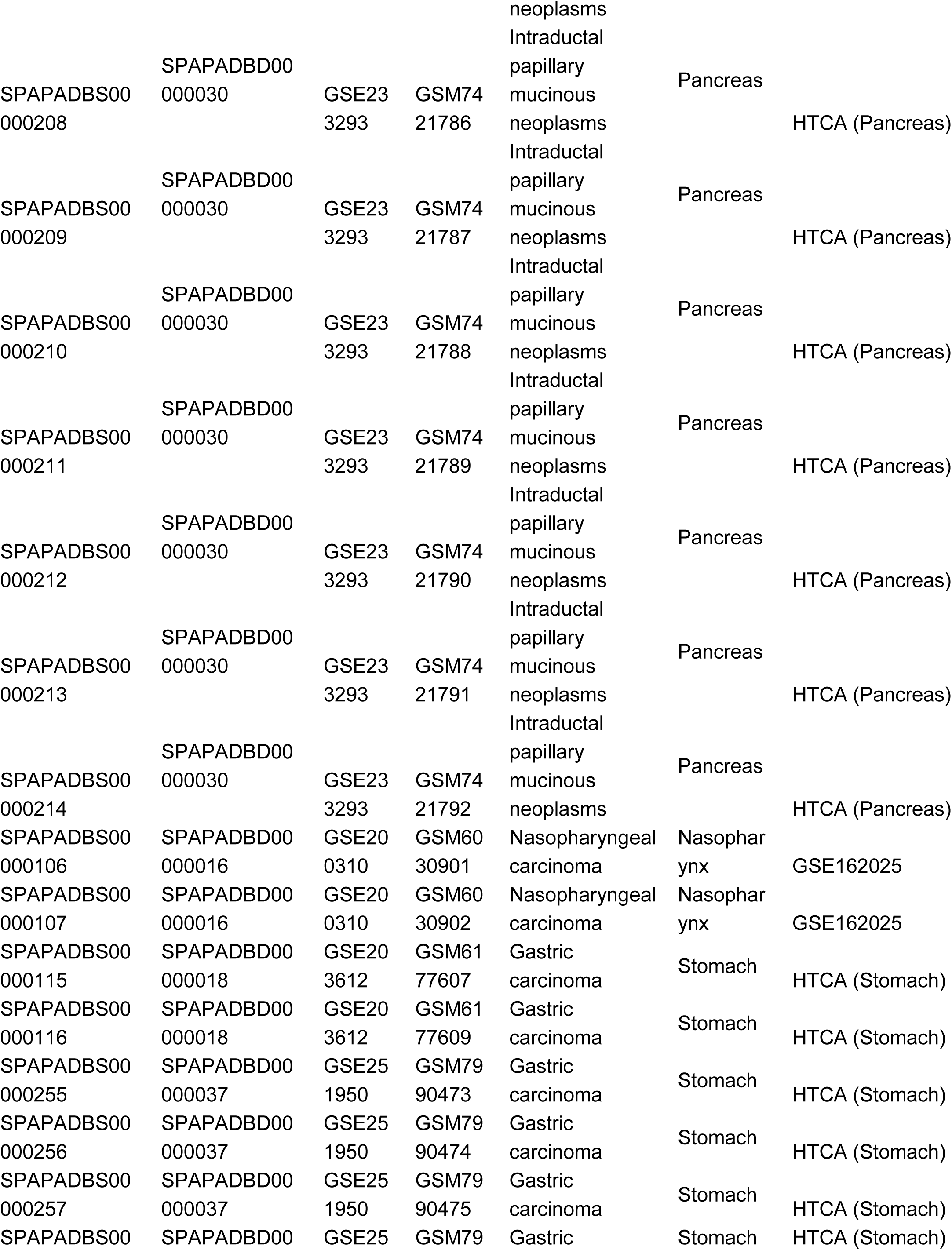
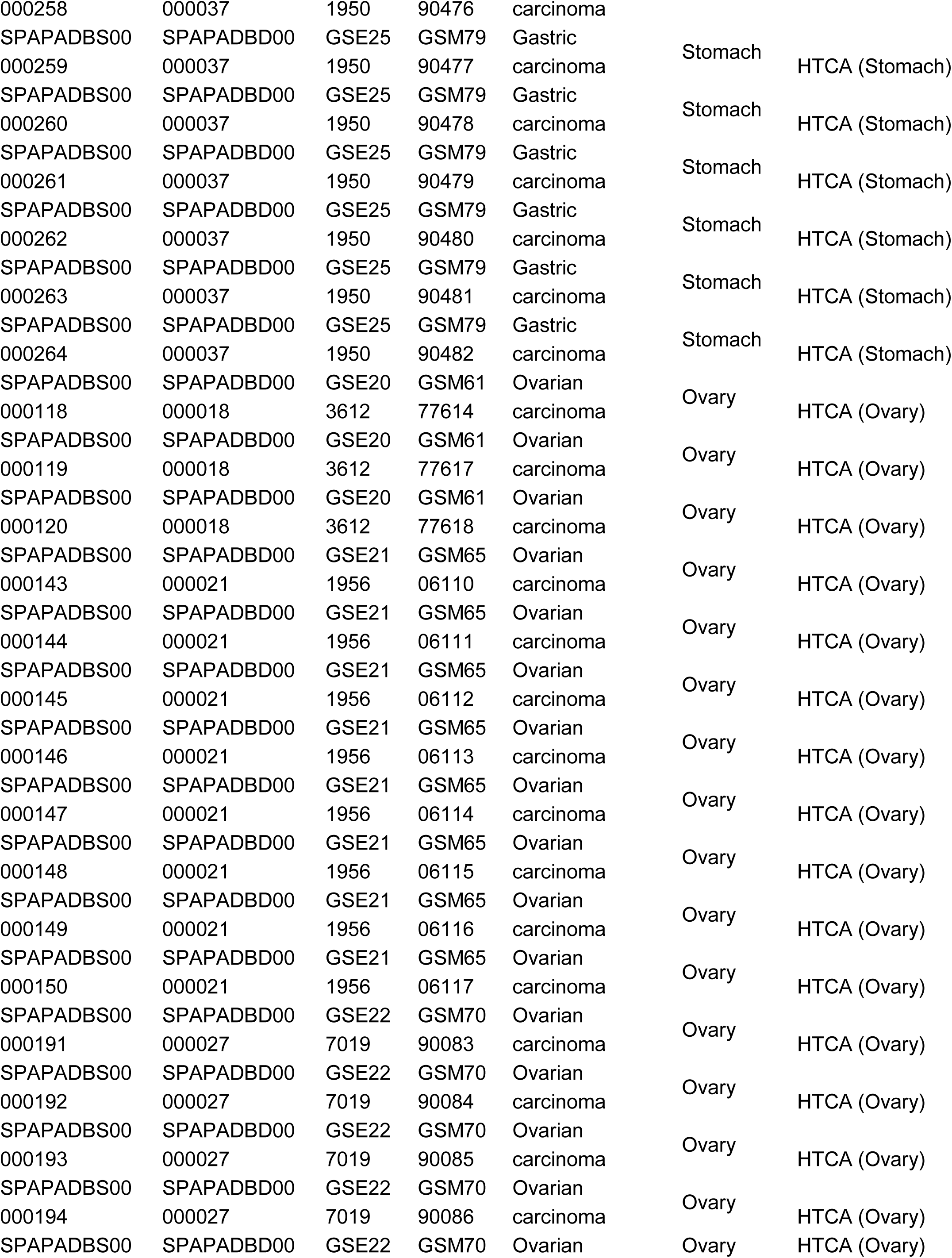
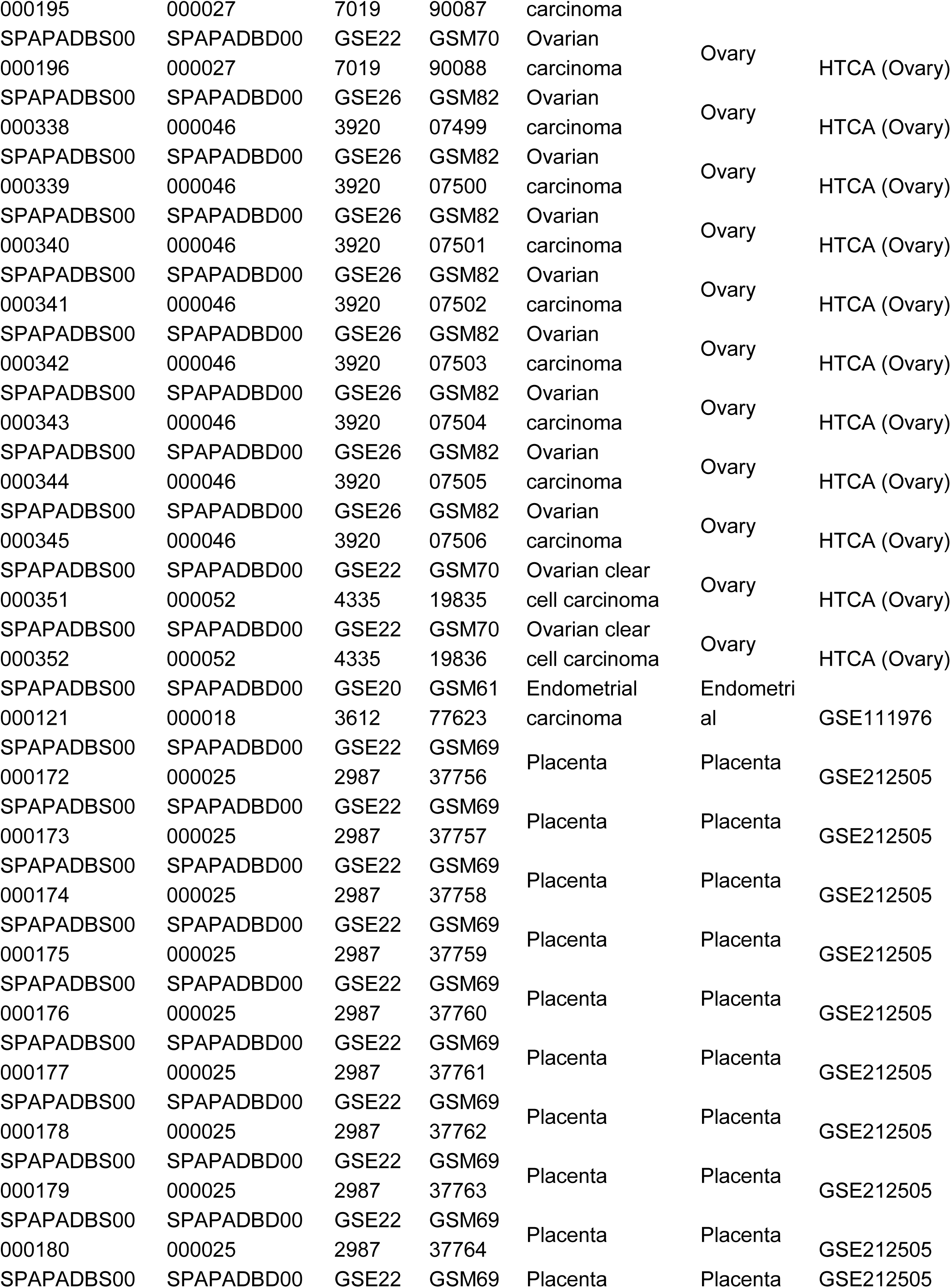
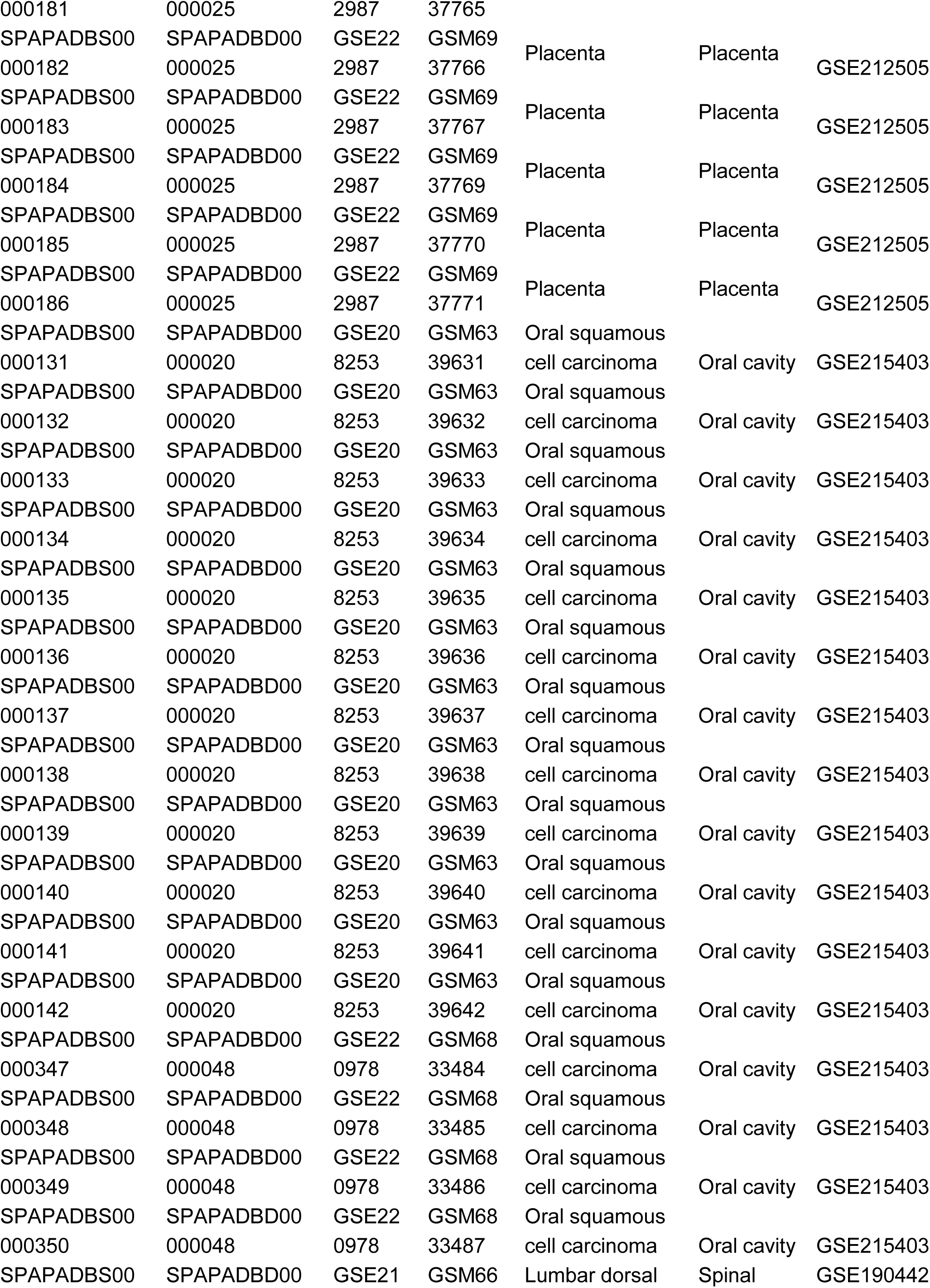
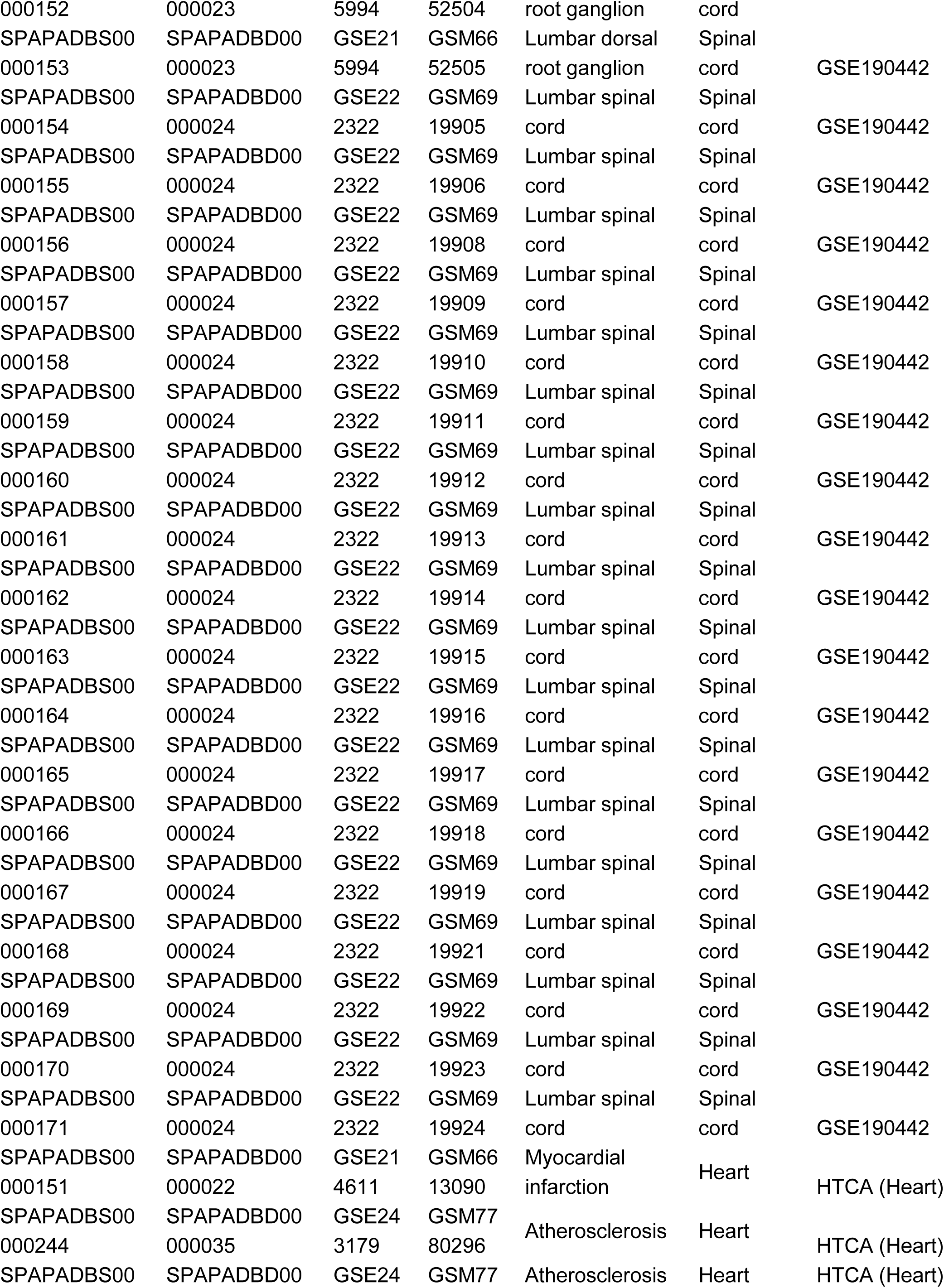
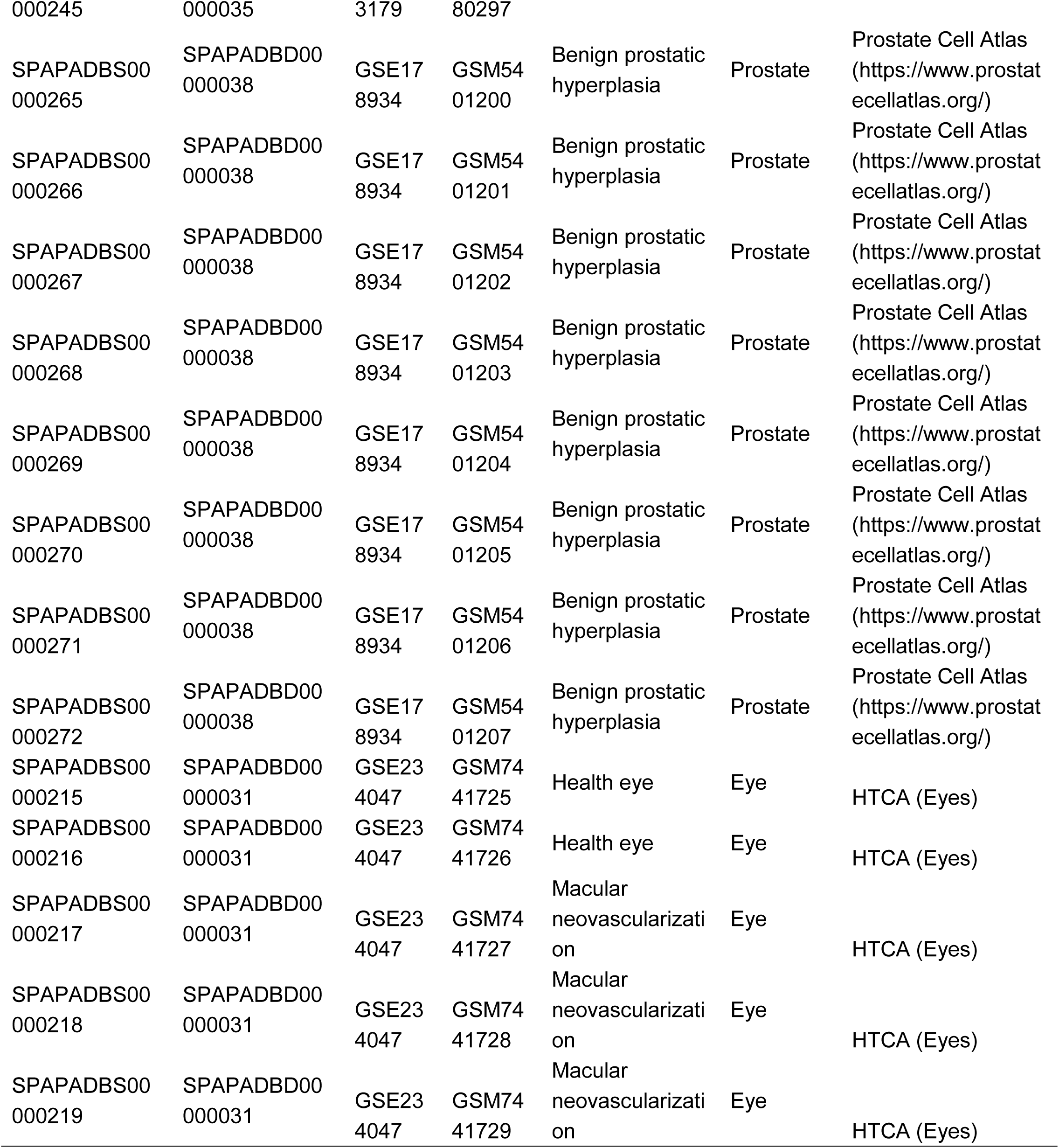
Detailed information on the reference datasets used for spatial cell deconvolution across various tissue organ types.

**Table S2.**
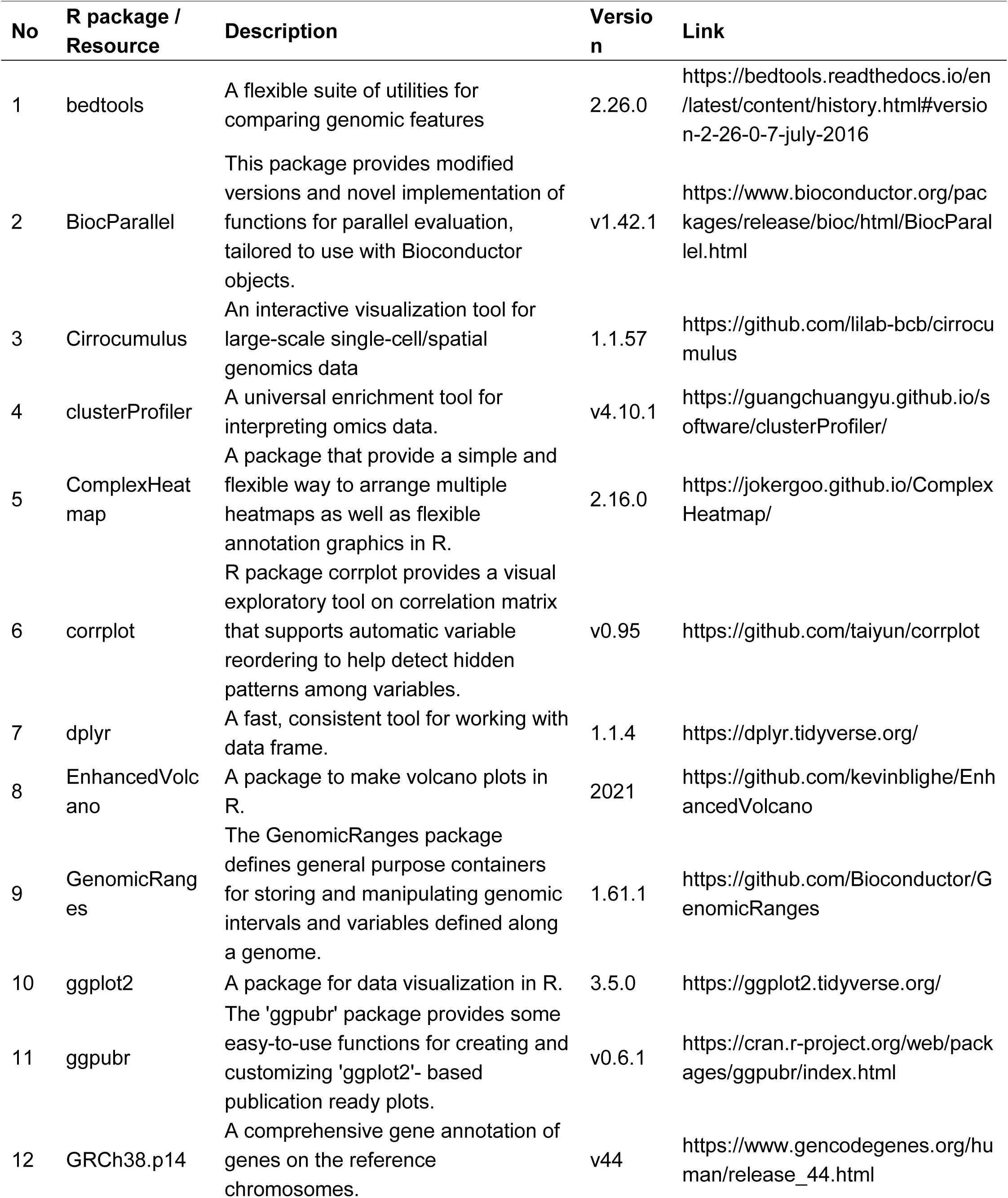
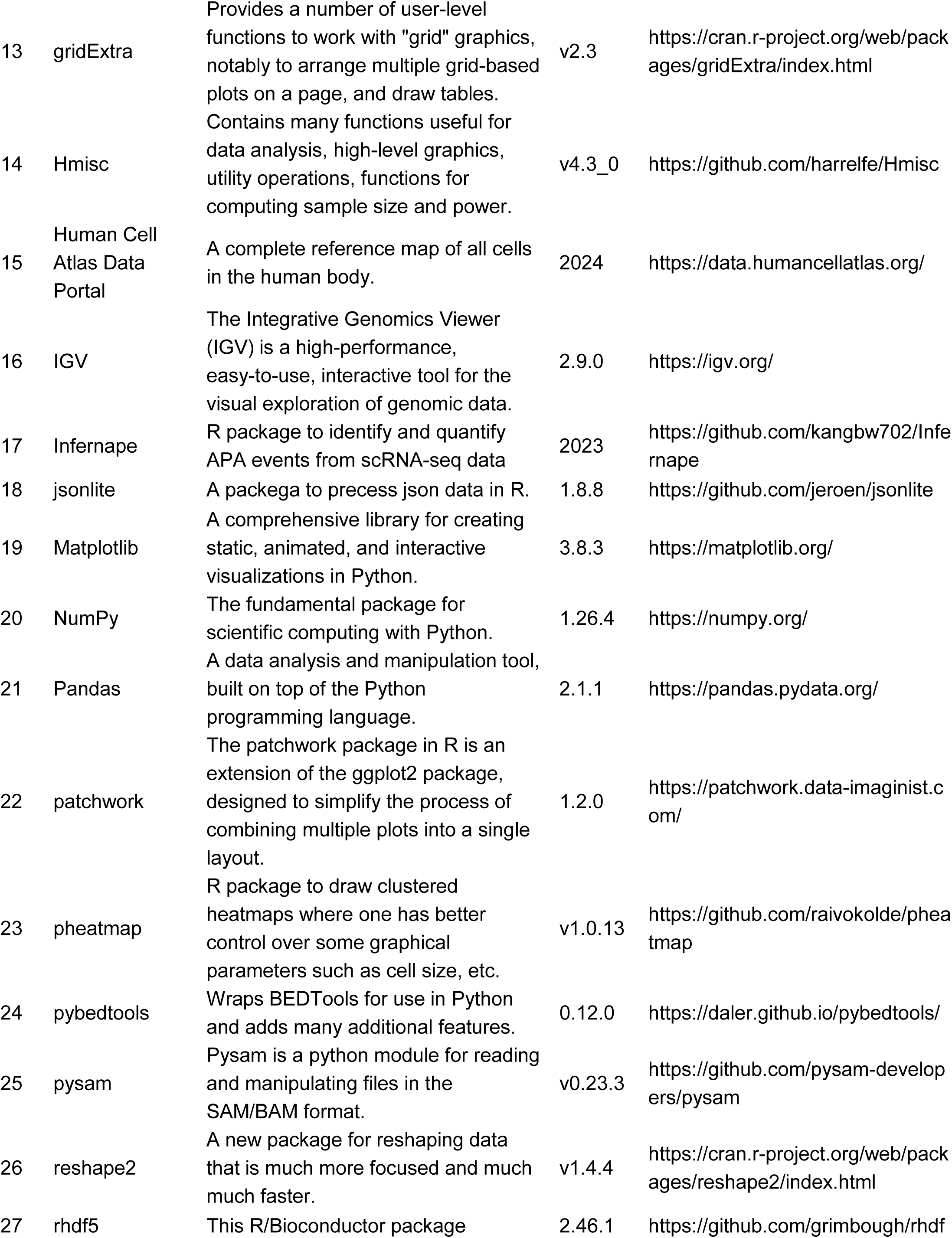
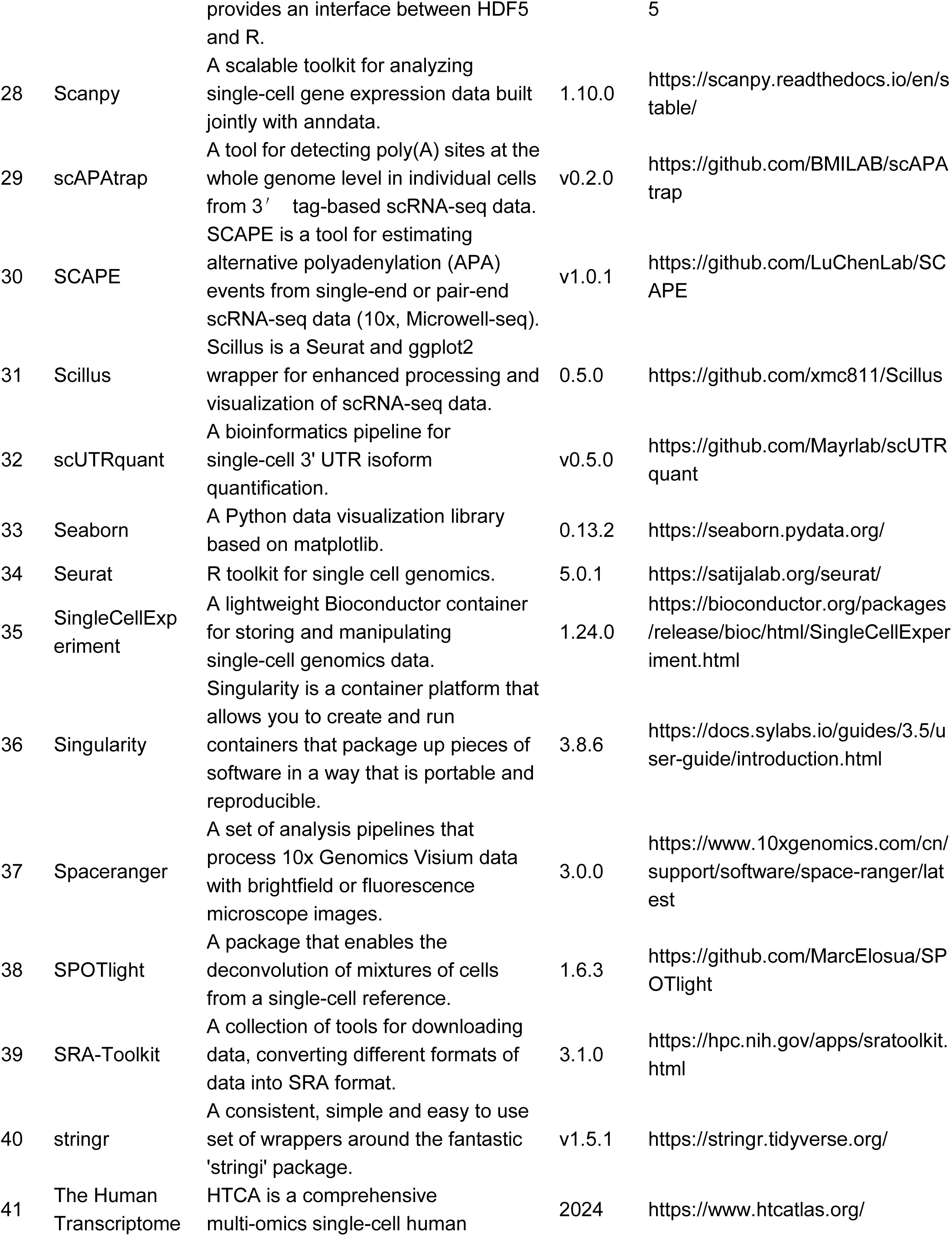
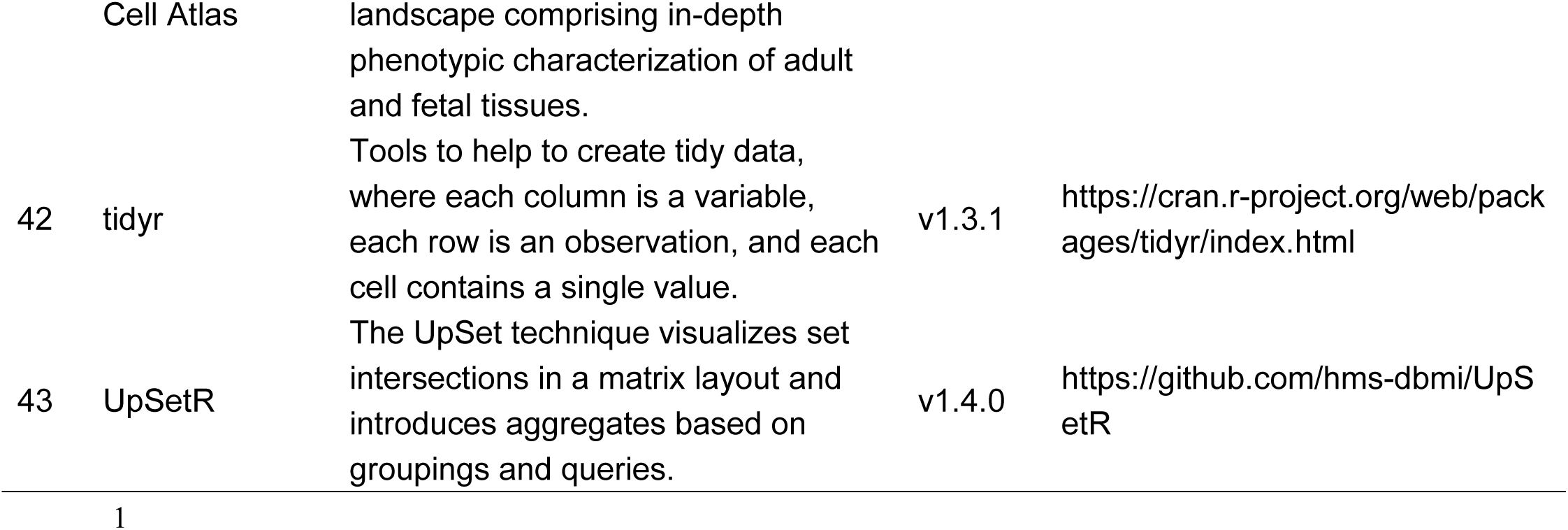
Details of key R packages and resources used for SpatialAPA and SpatialAPAdb.

